# An alternative mechanism of early nodal clustering and myelination onset in GABAergic neurons of the central nervous system

**DOI:** 10.1101/763573

**Authors:** Melina Thetiot, Sean A. Freeman, Thomas Roux, Anne-Laure Dubessy, Marie-Stéphane Aigrot, Quentin Rappeneau, François-Xavier Lejeune, Julien Tailleur, Nathalie Sol-Foulon, Catherine Lubetzki, Anne Desmazieres

## Abstract

In vertebrates, fast saltatory conduction along myelinated axons relies on the node of Ranvier. How nodes assemble on CNS neurons is not yet fully understood. We recently highlighted that clusters similar to nodes can form prior to myelin deposition in hippocampal GABAergic neurons and are associated with increased conduction velocity. Here, we used a live imaging approach to characterize the intrinsic mechanisms underlying the assembly of these early clusters. We first demonstrated that their components can partially pre-assemble prior to membrane targeting and determined the molecular motors involved in their trafficking. We then demonstrated the key role of the protein β2Nav for clustering initiation. We further unraveled the fate of these early clusters, by showing that they participate in node formation, but also have an unexpected role in guiding oligodendrocyte processes prior to myelin deposition. Altogether our results shed light on an alternative mechanism of nodal clustering and myelination onset.

## INTRODUCTION

The myelinated axon is regionally sub-compartmentalized into areas with differing conduction properties, as electrically insulated segments alternate with highly excitable, amyelinic domains, the nodes of Ranvier, thereby ensuring discrete yet complementary functions. Indeed, along myelinated axons, the nodes of Ranvier allow for rapid action potential regeneration and propagation by saltatory conduction (Huxley and Stampfli, 1949). The regeneration of electrical current at the nodes of Ranvier relies on the high concentration of voltage-gated sodium channels (Nav). The pore-forming Nav alpha-subunit (αNav) is complemented by the enrichment of cellular-adhesion molecules (CAMs) of the immunoglobulin domain (Ig) family, namely βNa_v_ subunits, which are associated with αNa_v_, as well as Neurofascin186 (Nfasc186), NrCAM and contactin, and of cytoskeletal scaffolding proteins such as AnkyrinG (AnkG) and ßIV-spectrin. All these proteins act in concert to target, traffic, and stabilize Nav at the nodes of Ranvier (Barry et al., 2014; Boyle et al., 2001; Çolakoğlu et al., 2014; Davis et al., 1996; Desmazieres et al., 2014; Dzhashiashvili et al., 2007; Eshed et al., 2005; Feinberg et al., 2010; Jenkins and Bennett, 2002; Jenkins et al., 2015; Namadurai et al., 2015; Sherman et al., 2005; Susuki et al., 2013; Yang et al., 2004; Zhang et al., 2012; Zonta et al., 2008). The mechanisms underlying the initiation of nodal clustering are well described in the peripheral nervous system (PNS) and rely on the tripartite interaction between axonal Nfasc186 and glial Gliomedin and NrCAM (Eshed et al., 2007, 2005; Feinberg et al., 2010; Zhang et al., 2012). The assembly mechanisms governing nodes of Ranvier clustering in the central nervous system (CNS) appear, however, to be more complex and involve an interplay of several overlapping and redundant mechanisms, ultimately converging on the fact that neuron-oligodendroglia interactions are necessary for CNS nodal assembly (Amor et al., 2017; Brivio et al., 2017; Dubessy et al., 2019; Freeman et al., 2015; Susuki et al., 2013).

The occurrence of evenly spaced clusters of nodal components along axons prior to myelination, initially reported on retinal ganglion cells (Kaplan et al., 2001, 1997) was also observed *in vitro* and *in vivo* on GABAergic hippocampal neurons, but not on pyramidal hippocampal neurons (Bonetto et al., 2019; Dubessy et al., 2019; Freeman et al., 2015). These early clusters, which share features with nodes of Ranvier (identical molecular composition and similar size), have a functional impact on neuronal physiology, as their presence correlates with increased axonal conduction of action potentials *in vitro* (Freeman et al., 2015). Interestingly most hippocampal Parvalbumin and Somatostatin expressing GABAergic neurons (i.e the neurons in which early clusters are detected) become myelinated *in vivo* (Stedehouder et al., 2017).

Furthermore, in multiple sclerosis, early nodal reclustering has also been observed in remyelinating plaques prior to myelin redeposition (Coman et al., 2006) following nodal disassembly associated with demyelination (Coman et al., 2006; Craner et al., 2004).

Many studies over the last two decades have focused on the neuron-glia interactions underlying CNS nodes of Ranvier assembly. Regarding the assembly of these early clusters, the role of oligodendroglial secreted factors is now established (Dubessy et al., 2019; Freeman et al., 2015; Kaplan et al., 2001, 1997); yet, in contrast, the role of the intrinsic neuronal mechanisms governing the trafficking, compartmentalization, and targeting of these early cluster proteins to the axolemma remains unexplored. More generally, the role of molecular motors in nodal protein transport is poorly understood. Recent works (Barry et al., 2014) reported that KIF5B, an anterograde microtubule molecular motor of the kinesin-1 superfamily, is able to transport Nav1.2 to the CNS nodes of Ranvier through associative linkage via AnkG. Whether other members of the kinesin-1 superfamily might participate in anterograde transport of proteins enriched at nodes has not been studied. Furthermore, the role of retrograde axonal cargo transport, which is mainly driven by the multisubunit dynein/dynactin protein complex along microtubules (Liu, 2017; Waterman-Storer et al., 1997), has yet to be explored with regards to the nodal component trafficking and targeting, as well as their clustering. In addition, whether the different proteins enriched at nodes are transported separately and assembled at the axonal membrane following targeting, or, in the contrary, associate early and are co-transported as a partially pre-assembled complex prior to cluster formation has yet to be elucidated.

Among the intrinsic cues at work in the clustering process, the possibility that several CAMs may orchestrate nodal assembly has been suggested by some reports. Nfasc186 has been described as the nodal clustering initiator on the axonal membrane in the PNS (Eshed et al., 2005; Feinberg et al., 2010; Zhang et al., 2012), while sodium channels subunits β1Nav and β2Nav have been shown to play critical roles in modulating the membrane density and the function of αNav (Chen et al., 2002; Kazen-Gillespie et al., 2000; O’Malley and Isom, 2015; Patton et al., 1994).

Furthermore, the relationship between the early clusters and the nodes of Ranvier has not been studied. The question as to whether these clusters are destined to disappear at the time of myelination, or persist and participate in the formation of nodes of Ranvier is open.

Here, by using live time-lapse imaging of fluorescently tagged nodal markers along axons in mixed hippocampal cell cultures, we address the mechanisms underlying early cluster protein trafficking and assembly, as well as the fate of these clusters. We show that kinesin-1 members KIF5A and C, as well as the dynein/dynactin complex, play a major role in their proteic components trafficking and clustering, and that a partial pre-assembly of the early cluster complex occurs prior to axonal membrane targeting. Using loss-of-function and overexpression experiments, we further provide evidence that β2Nav is a key player driving the clustering process. We finally demonstrate that early clusters not only persist, becoming heminodes when myelin deposition occurs, but can also act as localization signals guiding the deposition of myelin itself. Taken together these results unravel a key role of these early clusters and suggest an alternative mechanism of myelination onset and nodes of Ranvier formation within GABAergic neurons in the CNS.

## RESULTS

### Early cluster CAM trafficking is compatible with fast axonal transport characteristics

To address the mechanisms of early cluster protein trafficking, we focused on β1 and β2Na_v_ isoforms together with Nfasc186, as previous reports suggested they were the CAMs most likely associated with the early steps of nodal structures clustering (for review, Griggs et al., 2017; Winters and Isom, 2016).

We first established that fluorescently tagged and endogenous proteins were similarly localized in neurons, suggesting that their transport and proper targeting to the axonal membrane was not impaired (Fig. S1 A and not shown).

We followed the movement of the tagged proteins along the axon downstream of the AIS by performing time-lapse videomicroscopy at 17 days *in vitro* (DIV), at a time point when early clusters are actively forming (Freeman et al., 2015). The total number of puncta per 100 μm of axon per 50 s was higher for our proteins of interest compared to the control tag-only protein mCherry (which was potentially trapped unspecifically in some vesicles or accumulated in the cytoplasm), suggesting the studied proteins were specifically transported along the axon (Fig. S1 C). Punctate structures can be classified as moving (anterograde, retrograde, bidirectional) or stationary, and the tagged proteins of interest were mostly associated to dynamic puncta (Fig. S1 B and D). Transport characteristics were compatible with fast axonal transport associated to classical molecular motors such as kinesins and the dynein/dynactin-1 complex, respectively (for review, Maday et al., 2014).

### Early cluster proteins are partially co-transported along the axon, suggesting a pre-assembly of the protein complex prior to membrane targeting

Because we observed similar trafficking characteristics for the different early cluster markers, we then asked whether they could be co-transported along the axon.

To address this question, we co-expressed the early cluster proteins tagged with different fluorescent markers (GFP and mCherry) in mixed hippocampal cultures and monitored their axonal trafficking at 17 DIV with a dual-camera videomicroscopy system allowing for simultaneous streaming recordings of the two markers (Fig. 1 A and Movie S1). We performed co-expression of β2Na_v_-GFP together with APP-mCherry as an axonal protein control. APP has previously been described to be enriched at the CNS nodes of Ranvier in the late stage of their formation, after the initial Na_v_ channel clustering (Xu et al., 2014), thus making it unlikely to be co-transported with early-clustering nodal markers such as β2Na_v_-GFP.

**Figure 1.**
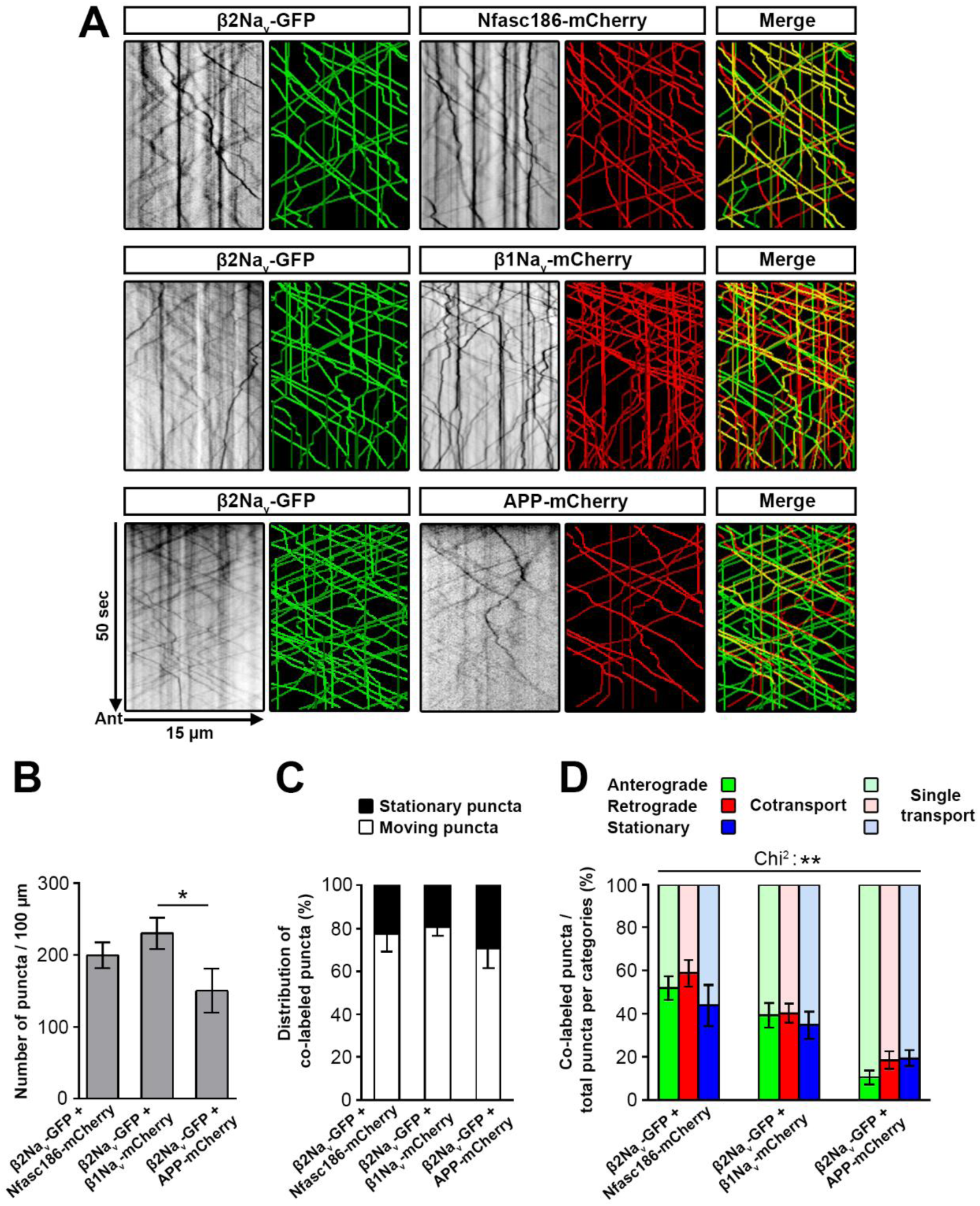
CAM proteins are partly co-transported in hippocampal neurons. **(A)** Co-expression of β2Na_v_-GFP and Nfasc186-mCherry, β1Na_v_-mCherry or APP-mCherry in hippocampal neurons at 17 DIV. Yellow tracks indicate axonal co-transport while tracks in red (mCherry) or green (GFP) show individual transport for these markers at 17 DIV. Mean number of puncta along the axon **(B)** and their distribution as moving or stationary structures **(C). (D)** Distribution of co-labeled puncta per categories. Data are mean ± SEM; n=17 to 22 neurons per condition from n=3 independent experiments. One-way analysis of variance with Tukey’s post-hoc test (**B**; **C**, *, p< 0.05; **, p < 0.01; and ***, p < 0.001). The difference of distribution between proteins across all types of movement **(D)** was assessed using Chi-squared test (see Table S2 for exact *p*-values).

We first quantified the total number of structures observed per 100 μm for each condition tested, which was slightly but significantly lower in the control condition (Fig. 1 B).

By overlapping the kymographs obtained for GFP-tagged (green) and mCherry-tagged (red) proteins, we observed in all tested conditions the existence of single-labeled (red or green trajectories) and co-labeled (yellow trajectories) puncta (Fig. 1 A). The vast majority of co-labeled structures were moving puncta, with only 19 to 30% of co-labeled structures being stationary (Fig. 1 C; compared to 83% of non-moving co-labeled puncta in a GFP/mCherry tag-only control condition, data not shown).

We then quantified the percentage of co-labeled structures compared to the total number of structures within each category (Fig. 1 D, anterograde, in green, retrograde, in red, and stationary, in blue). Interestingly, an important part of anterograde puncta were co-labeled for the CAMs (54% for β2Na_v_-GFP and Nfasc186-mCherry or 40% for β2Na_v_-GFP and β1Na_v_-mCherry), while this was rare in the control condition (11% for β2Na_v_-GFP and APP-mCherry, *p*-values < 0.0001). This suggests a specific anterograde co-transport of early cluster CAMs, likely reflecting a partial pre-assembly of the cluster complex prior to targeting at the axonal membrane.

### KIF5A and C act in synergy for the anterograde axonal transport of early cluster markers

KIF5B, one of the three members of the kinesin-1 family, was recently described to be implicated in Na_v_1.2 axonal transport (Barry et al., 2014). To address whether the neuronal kinesin-1 family members KIF5A and C could play a role in addition to KIF5B, we performed a knockdown of these proteins by specific miRNA expression, as single (KIF5A, Fig. S2 A or KIF5C, Fig. S2 B) or double knockdown (chained miRNA KIF5AC, Fig. S2 C and D). This approach led to a strong reduction of KIF5A (Fig. S2 A) and C expression (Fig. S2 B and D), though the loss of expression is only partial for KIF5A in the double knock-down context (Fig. S2 C and E; 42% reduction of expression as quantified by western-blot), which is classically observed for chained miRNA. In the double knockdown context, we further observed a slight increase in KIF5B expression (17%; Fig. S2 F), which suggests some compensatory expression may occur.

As illustrated for Nfasc186-mCherry at 17 DIV, the total number of puncta was not impaired when one single molecular motor was targeted (Fig. S3 A and B). However, we observed a significant 25% decrease in anterograde puncta and a 1.8-fold increase of stationary structures in KIF5A knock-down compared to the miRNA control condition (Fig. S3 C). In the KIF5C knockdown condition, we observed a significant 43% decrease of anterograde vesicles and a 1.9-fold increase of stationary structures compared to the control condition (Fig. S3 D).

These defects were strongly reinforced when performing the double KIF5AC knockdown, with a significant decrease of 44% in the total number of axonal fluorescent puncta per 100 µm together with a change in the distribution of the remaining structures (55% decrease of the anterograde vesicles fraction and a 2.3-fold increase of the stationary puncta fraction) in the KIF5AC knockdown compared to control condition (Fig. S3 E). Given the decrease of the total number of puncta and the reduction of the anterograde fraction for the remaining puncta, there is a 75% decrease of the number of anterograde vesicles per 100 μm in the axon in absence of both KIF5A and C, suggesting Nfasc186 axonal anterograde transport mainly relies on these molecular motors (Fig. 2 A and B).

**Figure 2.**
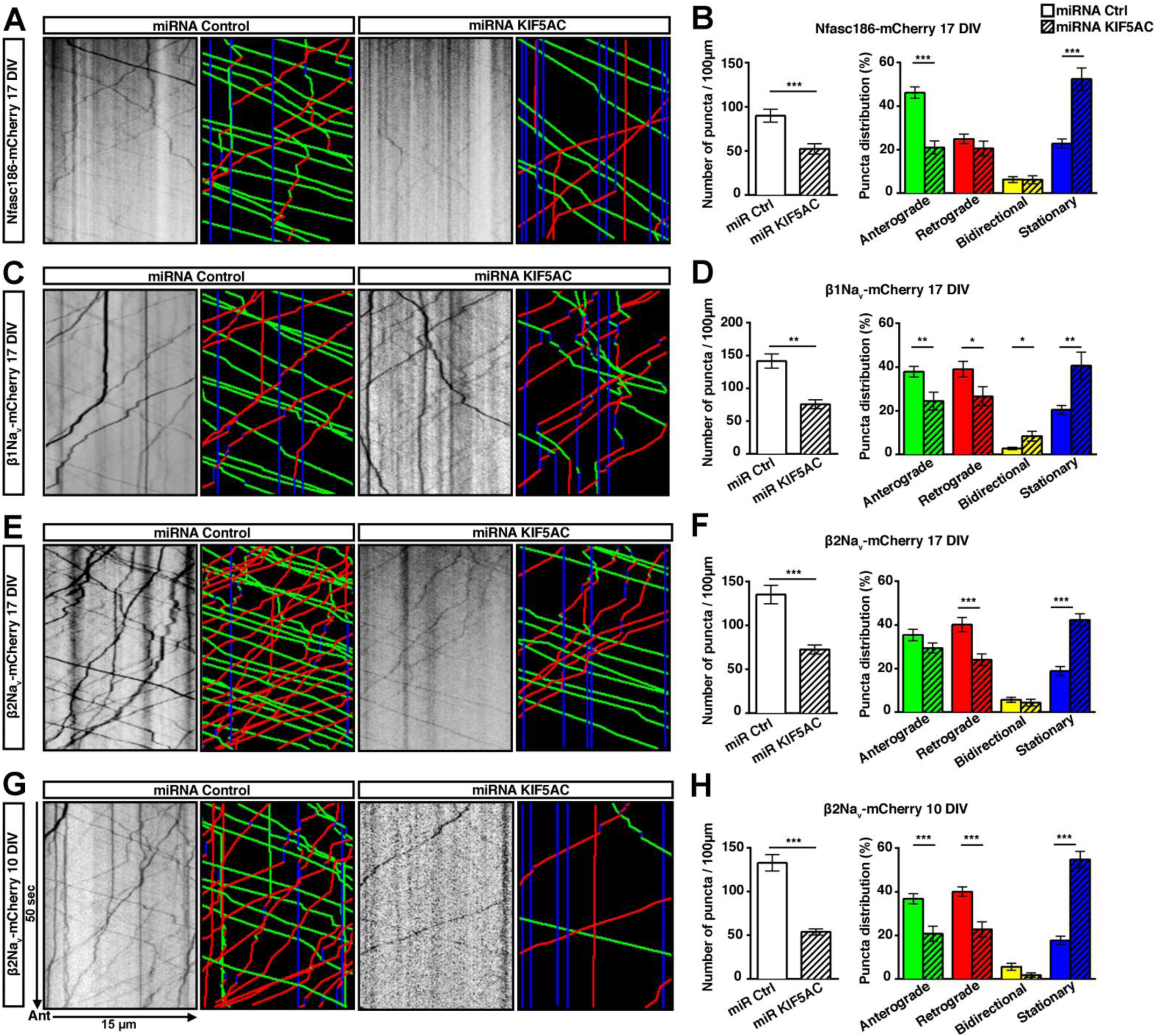
KIF5A and C participate in axonal transport of early cluster CAMs. Hippocampal mixed cultures were co-transfected with control or KIF5AC miRNA together with Nfasc186-mCherry **(A, B)**, β1Na_v_-mCherry **(C, D)** or β2Na_v_-mCherry (17 DIV: **E**, **F**; 10 DIV: **G**, **H**) expressing constructs. **(A, C, E, G)** Kymographs and corresponding tracks. **(B, D, F, H)** Mean number of puncta and distribution analysis of nodal proteins in control or KIF5AC double knockdown. Data are mean ± SEM; n=20 to 24 neurons per condition from n=3 independent experiments. Student’s two-tailed unpaired t-test (*, p < 0.05; **, p < 0.01; and ***, p < 0.001). The difference of puncta distribution between proteins across all types of movement was assessed using Chi-squared test (see Table S2 for exact *p*-values).

Similar results were obtained for β1Na_v_-mCherry (Fig. 2 C and D), with a 47% reduction of fluorescent vesicles in the axon, associated to a 36% decrease in the fraction of anterograde vesicles and a 1.9-fold increase in the fraction of stationary structures. Together, these data show a 66% decrease of the number of anterograde vesicles per 100 μm in the axon in the absence of KIF5A and C. Interestingly, β1Na_v_-mCherry retrograde transport was also significantly affected, with a 27% decrease in the knockdown condition.

Different results were obtained for β2Na_v_-mCherry (Fig. 2 E and F), suggesting its anterograde transport was less dependent on KIF5A and C than the other nodal CAMs at 17 DIV. Indeed, although we observed a significant 40% decrease in retrograde transport and the rate of stationary puncta increased by 2.2-fold, anterograde transport was mostly unaffected compared to the control condition at this time point (17% decrease, *p*-value: 0.0934). However, we observed a significant reduction in the number of total axonal puncta for β2Na_v_-mCherry (46% decrease), which suggested an impact of the loss of KIF5A and C on the anterograde flux of β2Na_v_-mCherry prior to 17 DIV.

To confirm this hypothesis, we analyzed β2Na_v_-mCherry trafficking at an earlier time point in the absence of KIF5A and C (10 DIV, Fig. 2 G and H). We observed a 60% decrease of total number of puncta, associated with reduced anterograde and retrograde transport (44% and 43%, respectively), as well as an increase in the fraction of stationary structures (3.1-fold). KIF5A and C suppression at 10 DIV thus resulted in a 77% decrease in the number of β2Na_v_-mCherry anterograde vesicles per 100 μm, confirming the early requirement of these motors for β2Na_v_-mCherry anterograde transport. These results suggest a different temporal regulation of axonal transport of β2Na_v_ compared to the other CAMs, with a switch of anterograde motors or the implementation of specific compensatory mechanisms for this protein at later time points.

In order to confirm that these results were specific to early cluster CAM trafficking and did not reflect a global alteration of axonal trafficking, we performed further axonal trafficking studies of lysotracker and synaptophysin-DsRed (Fig. S3 F and G respectively), as well as of APP-mCherry (which is known to be transported at least in part through the KIF5 family, Fig. S3 H and I), in the context of KIF5A and C double knock-down. No clear alteration of the global trafficking was observed in GABAergic neurons transfected with KIF5AC miRNA compared to control miRNA. In particular, there was a similar number of anterograde puncta in the case of APP-mCherry, when combining the total number of puncta per 100 μm and the puncta distribution (Fig. S3 I).

### Loss of dynactin-1, a protein of the dynein complex, impairs axonal transport of nodal proteins

To investigate the role of the dynein/dynactin-1 motor complex, which is classically associated to retrograde axonal transport, we performed a miRNA-mediated knockdown of dynactin-1/p150glued (Fig. S4 A, 17 DIV and S4 B, 21 DIV), which is required for dynein function by participating in the long-range stabilization of the complex along microtubules (Schroer, 2004; Tokito et al., 1996; Urnavicius et al., 2015).

Interestingly, dynactin-1 loss of function was associated with a reduction of the mean number of vesicles per 100 µm in the axon for Nfasc186-mCherry (Fig. 3 A and B) and β1Na_v_-mCherry (Fig. 3 C and D) at 17 DIV (39% and 21% respectively). In both cases, the fraction of retrograde transport was significantly decreased by 34%, alongside a significant 1.8-fold and 1.7 increase respectively in stationary puncta, while anterograde transport trended towards a reduction. This confirms a role of the dynein/dynactin-1 complex for the retrograde transport of early cluster markers, but also suggests a more general effect on their axonal trafficking.

**Figure 3.**
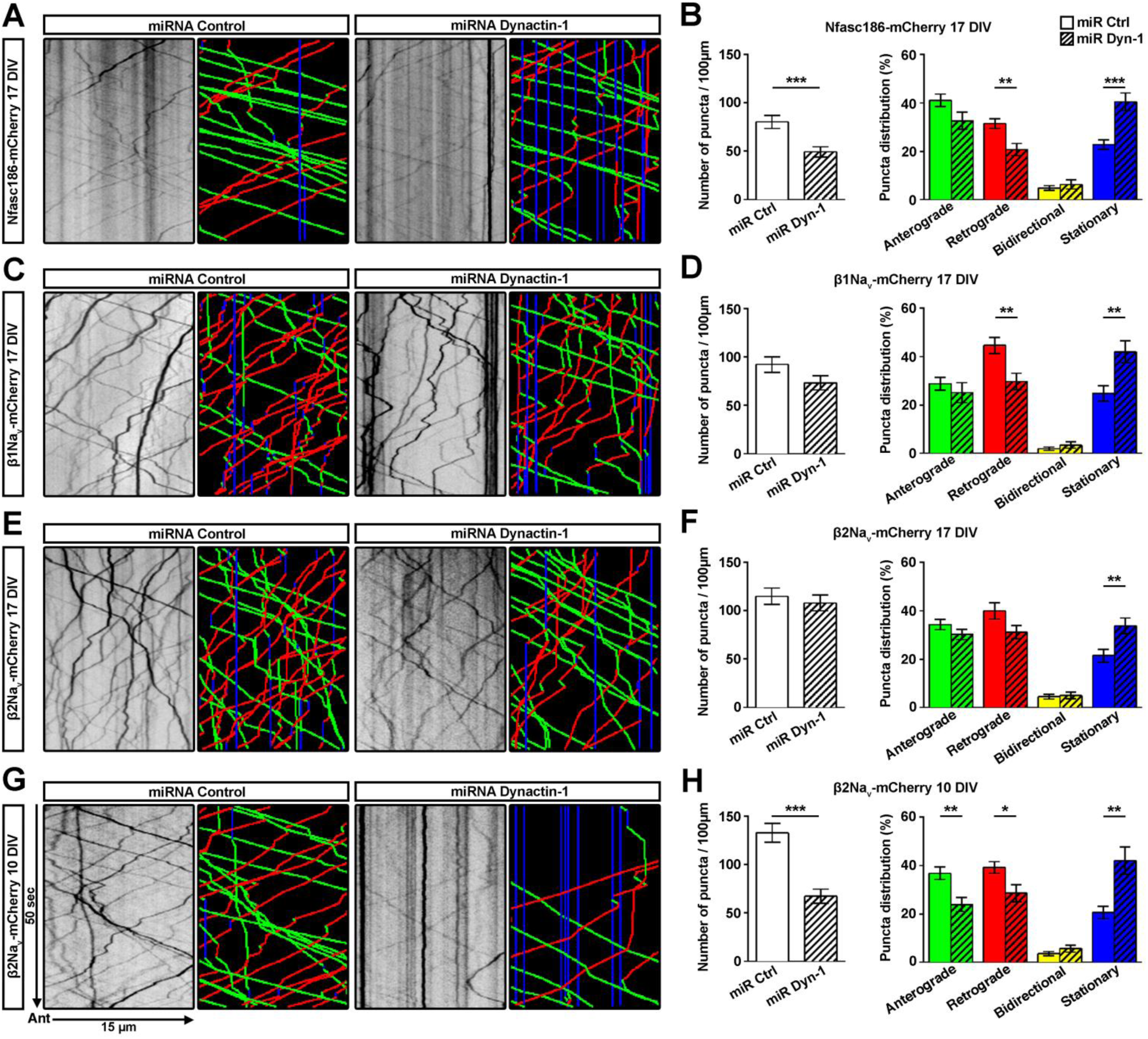
Dynactin-1 loss of function impairs CAMs transport. Hippocampal mixed cultures were co-transfected with control or Dynactin-1 miRNA together with Nfasc186-mCherry **(A, B)**, β1Na_v_-mCherry **(C, D)** or β2Na_v_-mCherry (17 DIV: **E**, **F**; 10 DIV: **G**, **H**) expressing constructs. **(A, C, E, G)** Kymographs and corresponding tracks. **(B, D, F, H)** Mean number of nodal CAMs puncta and their distribution in control or Dynactin-1 knockdown. Data are mean ± SEM; n=18 to 25 neurons per condition from n=3 independent experiments. Student’s two-tailed unpaired t-test (*, p < 0.05; **, p < 0.01; and ***, p < 0.001). The difference of puncta distribution between proteins across all types of movement was assessed using Chi-squared test (see Table S2 for exact *p*-values).

However, similarly to the KIF5AC double knockdown, β2Na_v_-mCherry was less affected by dynactin-1 loss of function than the other CAMs studied at 17 DIV (Fig. 3 E and F), with only a slight, statistically non-significant reduction of the total number of structures per 100 μm of axon and a 22% statistically non-significant decrease for retrograde vesicles (*p*-value: 0.0526). The only significant change in the absence of dynactin-1 at this time point was a 1.6-fold increase in stationary puncta. In contrast, when performing the dynactin-1 knock-down study at 10 DIV (Fig. 3 G and H), β2Na_v_ was profoundly affected, with a 50% reduction of the total axonal puncta. The distribution of remaining structures was similarly modified, with a significant decrease in anterograde and retrograde vesicles (35% and 28% respectively) and a 2-fold increase in stationary structures.

Taken together, these results show that both kinesin-1 family members KIF5A and C, as well as dynactin-1, are necessary for nodal protein trafficking prior to nodal clustering. In addition, they suggest that β2Na_v_ transport is differently regulated compared to β1Na_v_ and Nfasc186.

### Loss of kinesin-1 motors and dynactin-1 impairs early cluster assembly

Having characterized several molecular motors implicated in the axonal transport of early cluster CAMs, we then explored whether altering the transport of these proteins correlated with impaired early clustering (Fig. 4).

**Figure 4.**
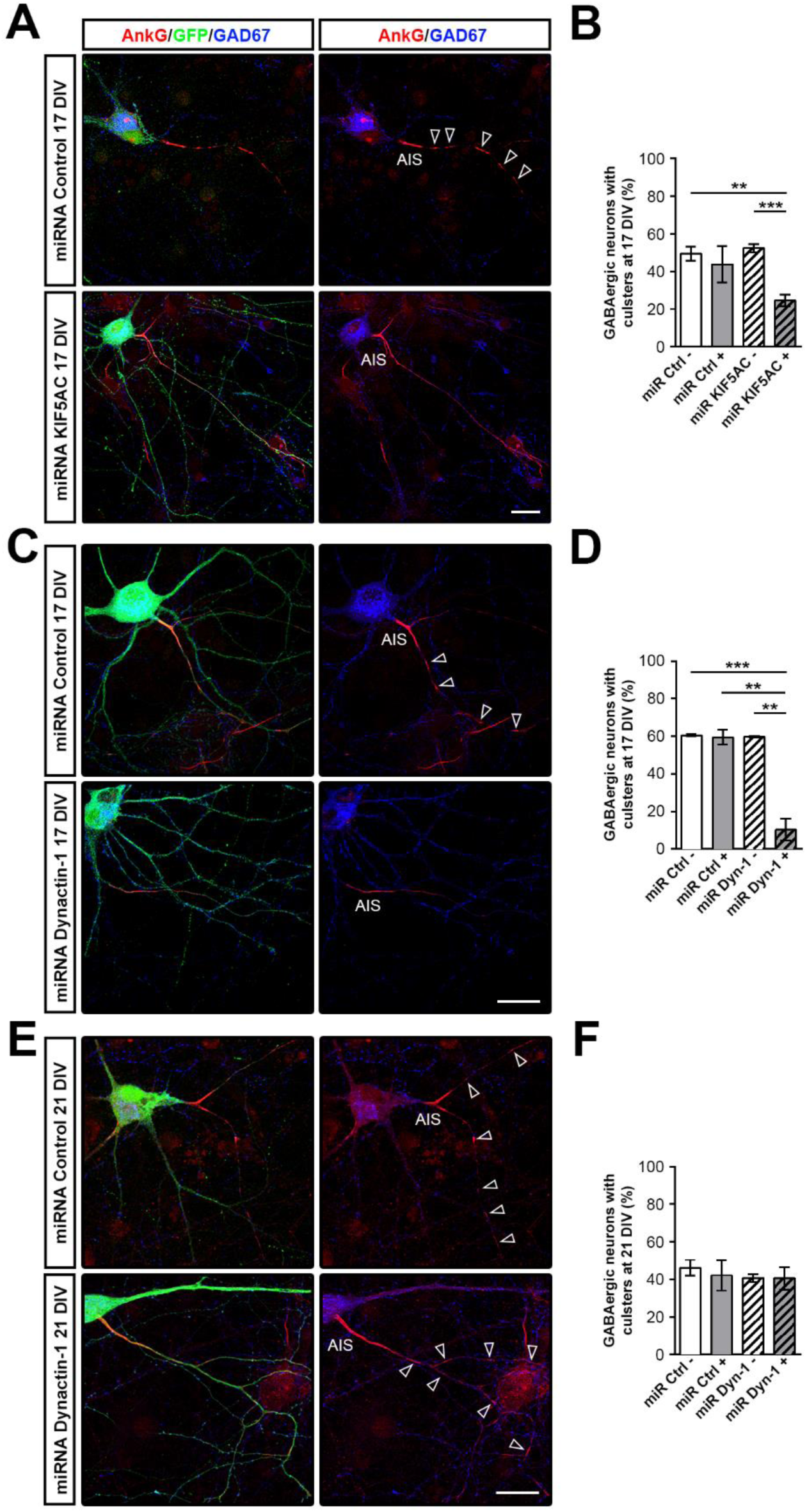
KIF5A, KIF5C and Dynactin-1 participate in early clusters assembly. **(A)** The presence of early clusters (AnkG, red, arrowhead) is impaired in GABAergic neurons (GAD67, blue) expressing KIF5AC miRNA (GFP, green) compared to control condition (GFP, green) at 17 DIV, in contrast to the axon initial segment (AIS) which is preserved. Nodal marker expression (AnkG, red) becomes diffuse along the axon in KIF5AC miRNA condition. Scale bars, 25 μm. **(B)** Quantification of GABAergic neurons with early clusters at 17 DIV in non-transfected neurons (miR Ctrl −/miR KIF5AC-) compared to transfected neurons with miRNA Control (miR Ctrl +) or miRNA KIF5AC (miR KIF5AC +) expressing constructs. **(C)** Early clusters (AnkG, red, arrowheads) are absent at 17 DIV in most GABAergic neurons (GAD67, blue) expressing Dynactin-1 miRNA (GFP, green) compared to control condition (GFP, green). **(E)** Early clusters (AnkG, red, arrowheads) are present at 21 DIV. Scale bars, 20 μm. Quantification of GABAergic neurons with early clusters at 17 DIV **(D)** or 21 DIV **(F)** in non-transfected neurons (miR Ctrl - or miR Dyn-1 -) compared to transfected neurons (miR Ctrl + or miR Dyn-1 +). Mean ± SEM of 3 to 4 independent experiments. Student’s two-tailed unpaired t-test; *, p < 0.05; **, p < 0.01; and ***, p < 0.001 (see Table S2 for exact *p*-values).

We observed a significant reduction by nearly 2-fold of the percentage of hippocampal GABAergic neurons with early clusters for the miRNA KIF5AC GFP^+^ cells (25% ± 3) compared to control conditions (miRNA Ctrl GFP^−^: 49% ± 4; miRNA Ctrl GFP^+^: 44% ± 10; miRNA KIF5AC GFP^−^: 52 ± 2). This suggests that the double knockdown of KIF5A and C impacts the clustering, but that the remaining axonal transport preserves it in some neurons. In order to confirm the specificity of this alteration, we further performed loss of function for other kinesins known to be implicated in axonal transport (KIF3, KIF4 and KIF1A) through miRNA or negative dominant approaches (Fig. S5 A, B, and E (Xue et al., 2010)). In such conditions, we did not observe any clear clustering alteration compared to the control conditions (Fig. S5 C-F), confirming the predominant role of the KIF5 family in early clustering.

Furthermore, the loss of dynactin-1 had a major effect on the clustering, resulting in a 6-fold reduction of the percentage of GABAergic neurons with early clusters at 17 DIV (Fig. 4 C and D).

This reduction of clustering corresponded to a delay rather than a loss of clustering, as demonstrated by the detection of early clusters at later time points (21 DIV), with no significant difference in the percentage of miRNA Dyn-1 transfected GABAergic neurons compared to control conditions (Fig. 4 E and F).

The dynein/dynactin-1 complex has been described to participate in neuronal differentiation and survival through axonal retrograde transport of neurotrophic factors bound to their internalized receptor (for review, Howe and Mobley, 2005; Roberts et al., 2014). To assess whether the effects of dynactin-1 loss could be due to a delay in maturation of the targeted neuron, we performed immunostainings against phosphorylated neurofilaments using SMI31 antibody (data not shown), as well as against Neurofilament-M (NfM), which expression increases with maturation, at 14 DIV, when initiation of the clustering normally occurs (Fig. S4 C). We observed a strong reduction of their expression in the absence of dynactin-1 compared to the control conditions at 14 DIV in GABAergic neurons (though no clear difference was seen at 17 DIV, data not shown). This data suggests that the impact of dynactin-1 on the clustering could be partly linked to neuronal maturation, without excluding a direct effect in nodal protein transport.

Taken together, these results suggest that trafficking parameters linked with specific anterograde and retrograde molecular motors are required for early clustering. We then wondered whether some early cluster proteins themselves could play the role of organizers for this clustering.

### *β2Na*v accumulates prior to other CAMs at discrete sites distally to the last cluster formed

While performing the co-transport videomicroscopy study, we observed focal enrichments of β2Na_v_ along the axon, which was not seen for the other early cluster CAMs (Fig. 5A and Movie S2). This was confirmed with various combinations of immunostainings for the endogenous proteins at early stages of clustering (14 DIV and 17 DIV, Fig. 5 B-F) in non-transfected cultures. The clustering is sequential, from the axon initial segment (AIS) vicinity to the distal part of the axon (Freeman et al., 2015). We observed enrichments of β2Na_v_ distally to the last cluster, while Nfasc186, NrCAM and β1Na_v_ were not detected at such locations (Fig. 5 B, D, and F, respectively). In contrast, αNa_v_ and AnkG always colocalized with β2Na_v_ (Fig. 5 C and E, respectively). These data suggest an initial clustering of a complex of β2Na_v_, AnkG and αNa_v_ channels, while other CAMs are targeted subsequently. This hypothesis of a two-step mechanism is supported by live-imaging videos showing broad clusters becoming restricted with time (Movie S3), a feature we regularly observed.

**Figure 5.**
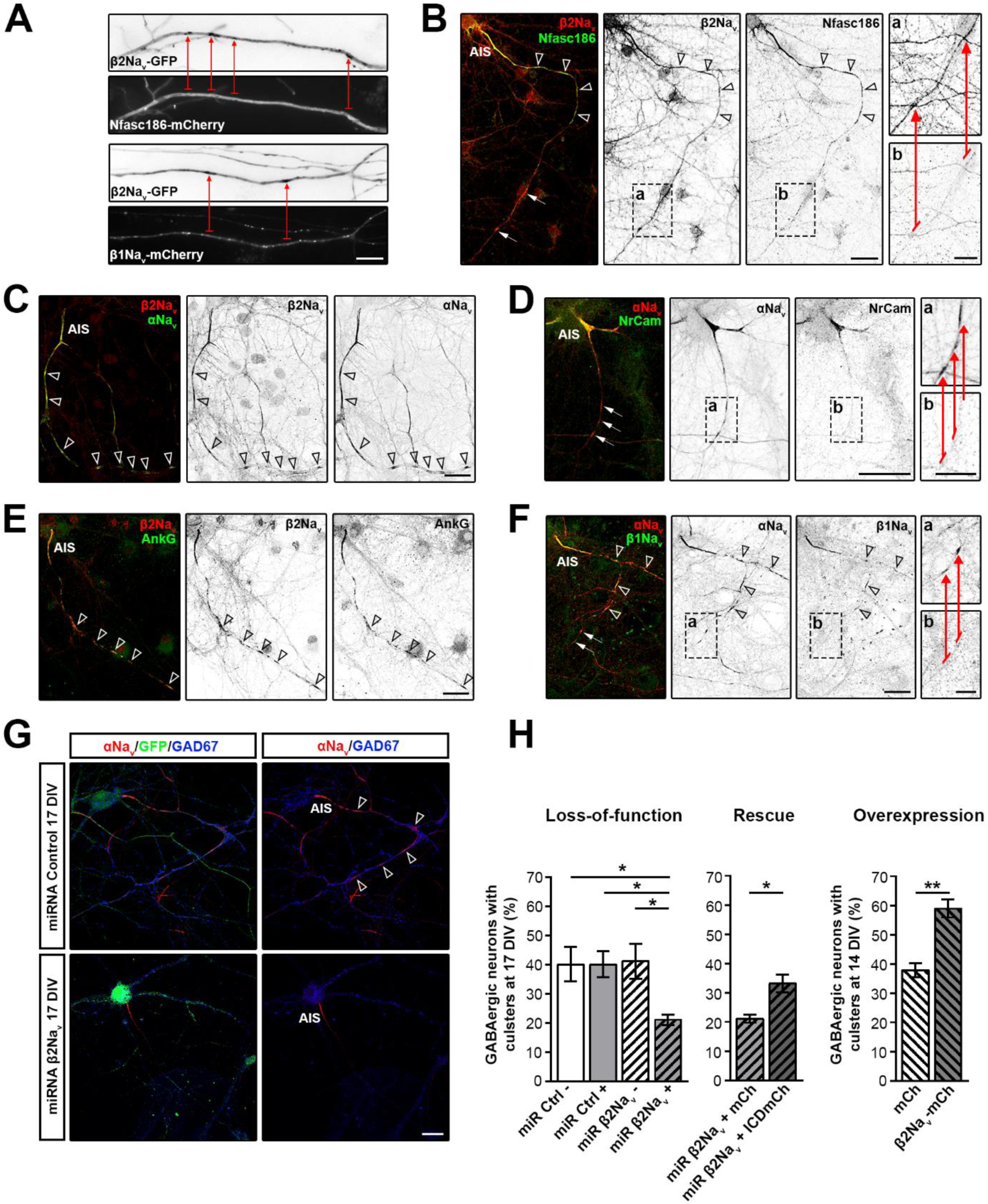
β2Na_v_ is a key player of early clusters initiation. **(A)** Early axonal accumulation of β2Na_v_-GFP (red arrows) in absence of Nfasc186-mCherry or β1Na_v_-mCherry (red arrows counterpart) at 17 DIV in live cells. **(B)** β2Na_v_ (red) is enriched distally to the last cluster formed (arrowheads), prior to Nfasc186 accumulation (green, arrows). β2Na_v_ clusters (in red) always colocalize with αNa_v_ and AnkG (in green, **C** and **E** respectively, arrowheads). At 17 DIV, some distal clusters express αNa_v_ (red) without NrCAM or β1Na_v_ (in green, **D** and **F**, respectively, arrows). **(G)** Loss of early clusters (αNa_v_, red, arrowheads) in GABAergic neurons (GAD67, blue) expressing β2Na_v_ miRNA (GFP, green) compared to control miRNA (GFP, green) at 17 DIV. AIS: axon initial segment. Scale bars, 5 µm **(A)**; 25µm **(B-G)**; 10 µm (**a** and **b** boxes). **(H)** Quantification of GABAergic neurons with clusters at 17 DIV in non-transfected neurons (miR Ctrl - or miR β2Na_v_ -) compared to transfected neurons (miR Ctrl + or miR β2Na_v_ +). At 17 DIV, GABAergic neurons transfected with the β2Na_v_ miRNA co-expressing mCherry (miR β2Na_v_ + mCh) or ICD-mCherry (miR β2Na_v_ + ICDmCh) show a partial rescue of clustering by ICD in absence of β2Na_v_. At 14 DIV, GABAergic neurons overexpressing β2Na_v_-mCh show an increase in GABAergic neurons with early clusters compared to control condition (overexpression of mCherry alone). Mean ± SEM of 3 independent experiments. Student’s two-tailed unpaired t-test; *, p < 0.05; **, p < 0.01; and ***, p < 0.001 (see Table S2 for exact *p*-values).

### Early clustering partly relies on *β2Na*v

In previous studies, we showed by miRNA-mediated knockdown that AnkG is required for early cluster assembly (Freeman et al., 2015). The early association of β2Na_v_ with AnkG and αNa_v_ channels in early clusters prompted us to address whether β2Na_v_ could be required for the initiation of Na_v_ channel clustering.

To address this question, we performed β2Na_v_ miRNA-mediated knock-down in primary cultures of rat hippocampal neurons (Fig. S6 A), and quantified its impact on early cluster formation.

We first quantified the percentage of transfected GABAergic neurons with clusters at 17 DIV in control and knockdown conditions (GAD^+^GFP^+^) and the percentage of non-transfected GABAergic neurons with clusters (GAD^+^GFP^−^) in the same cultures. While there was no significant reduction in the percentage of GABAergic neurons with clusters in GFP^+^ neurons compared to GFP^−^ neurons in the control condition (40% ± 6 vs 40% ± 5), we observed a strong reduction of GABAergic neurons with clusters following β2Na_v_ knockdown (GFP^+^: 21% ± 2 vs GFP^−^: 41% ± 6) (Fig. 5G and H), showing that β2Na_v_ is a key protein in early clustering.

β2Na_v_ plays a direct role in αNa_v_ targeting through its adhesion properties (Chen et al., 2002; Malhotra et al., 2002), but also participates in the regulation of αNa_v_ expression itself, through its intracellular part (ICD) (Kim et al., 2007, 2005). ICD is generated by cleavage of the β2Na_v_ targeted at the membrane and trafficked back to the nucleus, where it, in particular, regulates the transcription of the Na_v_1.1 isoform. In order to assess which of these functions of β2Na_v_ (or both) was required for early clustering, we generated a construct allowing the neuronal expression of ICD tagged with mCherry. We then coexpressed the miRNA β2Na_v_ together with ICD-mCherry, or mCherry as a control (Fig. S6 C).

ICD expression significantly rescues early clustering in the absence of β2Na_v_ at DIV17, with 33% ± 3 of GABAergic neurons co-transfected showing clusters, compared to 21% ± 1.4 for the miRNA β2Na_v_/mCherry condition (which is similar to the miRNA β2Na_v_ only condition), and to about 40% for control non transfected GABAergic cells (Fig. 5 H).

We then wondered whether β2Na_v_ could be sufficient to induce early clustering. To test this hypothesis, we overexpressed β2Na_v_-mCherry, or mCherry as a control, in neurons in mixed hippocampal cultures (Fig. S6 B). The percentage of GABAergic neurons expressing β2Na_v_-mCherry with clusters was significantly higher (59% ± 3), compared to the GABAergic neurons expressing mCherry with clusters (38% ± 2).

### Early clusters participate in heminodal formation

Multiple and complementary mechanisms have been suggested to participate in node of Ranvier clustering. However, the potential contribution of early clusters in the formation of mature nodes has never been addressed. To answer this key question, we performed live imaging between 20 and 23 DIV on mixed hippocampal cultures complemented with PLP-GFP mouse oligodendrocytes, using an antibody directed against a Neurofascin external epitope directly coupled to Alexa-594 to stain nodal structures. This model allows us to visualize the structures enriched in nodal proteins along the axon (heminodes, mature nodes and early clusters, in red) and their fate when myelination proceeds (oligodendrocytes and myelin, in green, Fig. 6).

**Figure 6.**
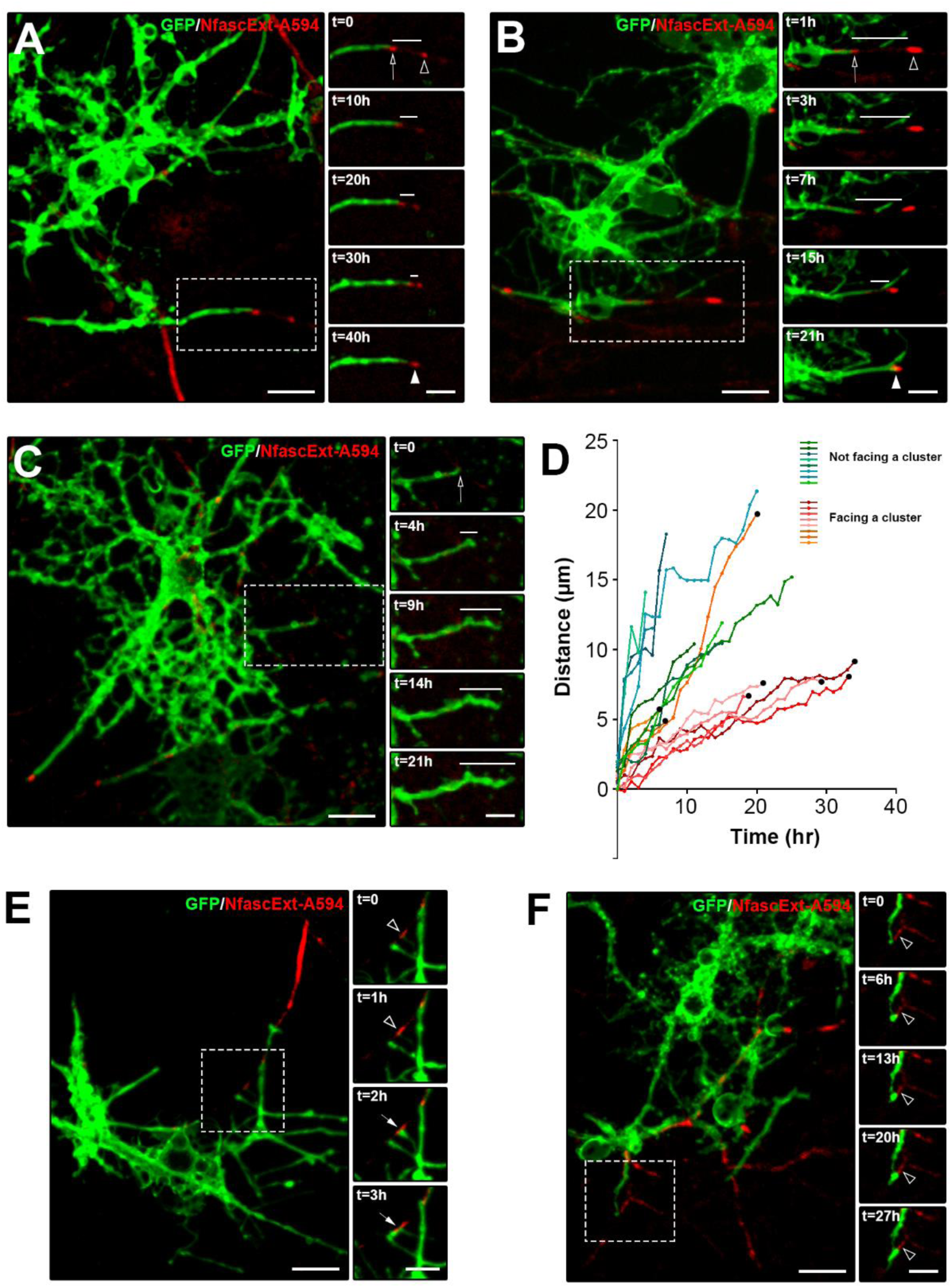
Live-imaging of early clusters fate during myelination *in vitro*. **(A-C; E and F)** For each live image, the boxed area is shown along time (right). Oligodendrocytes and myelin sheaths are visualized through GFP (green) and early clusters are revealed by live staining of the external epitope of Neurofascin (NfascExt-A594, red). **(A, B)** Fusion of a heminode (arrow) with an early cluster (arrowhead) during myelination (the line indicates the gap between the two structures at a given time). **(C)** Myelin growth along an axon without cluster at its vicinity (the line indicates myelin progression from the beginning (arrow) to the end of the imaging). **(D)** Distance covered by individual myelin sheaths along time. One color represents one individual myelin sheath with red-orange curves corresponding to individual myelin sheaths facing early clusters, while green-blue curves correspond to individual myelin sheaths without a cluster facing the progressing myelin tip. Black dots indicate the time point when a heminode and a cluster fuse. n=8 individual myelin sheaths for each condition (red/orange or blue/green curves). **(A)** is representative of the red pool of curves, while **(B)** shows the atypical high speed growth of myelin corresponding to the orange curve on **(D)**. **(E)** An early cluster (red, arrowheads) is contacted by an oligodendrocyte (green, white arrows), which then starts to wrap the axon at its vicinity **(E, F).** Scale bar, 15 µm; boxes, 10µm.

We observed that early clusters were not suppressed when myelination proceeds, as an early cluster disappearance was detected in only one out of the 18 movies analyzed. The early clusters rather persist and become part of the heminodes at the tip of the progressing myelin (Fig. 6 A and B, Movie S6; 10 out of 10 movies where the myelin and heminode reach the early cluster location). Interestingly, we further observed the existence of a second scenario: a contact occurs between an oligodendrocyte process and an early cluster, which is followed by the onset of myelin deposition in the “peri-cluster area”, the early cluster thus becoming a heminode (Fig. 6 E and F, Movies S4 and S5; 7 movies). Moreover, we did not observe initiation of myelination elsewhere along the axon of GABAergic neurons with early clusters (than at the close vicinity of the clusters). Early clusters can thus become heminodes through two different processes.

In addition, in the first scenario, when the myelin tip and the associated heminode were getting close to the early cluster, a diffusion of the heminode and/or the early cluster labeling was regularly evidenced (heminode or early cluster, 6/10 movies; both, 3/10 movies) before the myelin reaches the cluster. When diffusion is detected in only one of the two structures, it was in most cases the progressing heminodal structure (Fig. 6 B; Movie S7; 5/6 movies), rather than the early cluster (1/6), suggesting that the moving structure is more labile.

We further measured the distance covered along time by elongating myelin sheaths on axons with (Fig. 6 D, red and orange curves) or without early clusters (Fig. 6 C and D, green and blue curves). The cumulated distances covered along time were significantly reduced when myelin elongated at the vicinity of a cluster (mean speed of progression of 0.3 ± 0.1 μm/hr) compared to myelin elongation in the absence of an early cluster closeby (mean speed of progression of 1.1 ± 0.2 μm/hr, *p-*value: 0.0028, Mann-Whitney test). Furthermore, the mean speed of myelin elongation tended to decrease the closer the myelin tip was from the cluster (Fig. 6 D, red curves, from 0.3 ± 0.196 μm/hr, when the cluster was more than 5μm away, to 0.2 ± 0.03 μm/hr when it was less than 5 μm away, n=5 individual myelin sheaths, *p*-value: 0,056). This suggests that axonal structure changes might take place when the myelin tip reaches the early cluster area.

Taken together, this study shows for the first time that early clusters not only persist over time but that they are committed to participate in heminode formation (which will later result in nodes of Ranvier, see discussion), and may also act as a localization signal thereby guiding myelination onset.

## DISCUSSION

Here, we provide new insights into the dynamics of CNS early cluster protein trafficking and identify novel mechanisms leading to their clustering. Our data furthermore suggest that early clusters, i.e node-like clusters formed before myelination, participate in heminode formation (heminodes will then fuse to become nodes of Ranvier) and guidance of myelination initiation.

### Partial preassembly of the node-like complex prior to early cluster formation

By following the trafficking of different early cluster proteins in the same axon, our live imaging data show that significant proportions of β2Na_v_ and β1Na_v_, but also of β2Na_v_ and Nfasc186 are co-transported, in particular anterogradely, suggesting that the node-like complex partially assembles prior to clustering at the axonal membrane.

This partial pre-assembly of the early cluster complex could be due to a direct association of different node-like proteins when trafficking vesicles are formed following protein synthesis. Alternatively, it might result from an initial node-like complex assembly at the soma membrane prior to recycling through endocytosis and trafficking to the axon, as previously suggested (Fache et al., 2004; Lai and Jan, 2006; Wisco et al., 2003).

Retrograde co-transport could on the other hand participate in node-like marker reorganization (retrieval of excess proteins, recycling, or membrane targeting), as the literature suggests excitable axonal domain protein isoforms and density can be finely regulated (Bréchet et al., 2008; Dzhashiashvili et al., 2007; Jamann et al., 2018; Leterrier et al., 2011; Rasband, 2008; Zhang et al., 2012).

Of note, not all proteins were co-transported, as shown for APP, a late nodal component, which is mostly transported independently when node-like cluster form. Furthermore, a subpart of the node-like CAMs trafficked separately, which is compatible with the slight delay in enrichment at early clusters observed for some CAMs compared to β2Na_v_.

This suggests a finely tuned temporal regulation of node-like protein trafficking, with different transport strategies within the initial and late phase of the clustering.

### Kinesin-1 and dynein/dynactin-1 motors are involved in node-like protein trafficking and early clustering

In the PNS, the sequence of protein delivery and enrichment at the nodes has been partly deciphered (Zhang et al., 2012). As shown in cultured dorsal root ganglion neurons, sodium channel localization might depend on different mechanisms, relying on long-range transport of the protein by molecular motors, such as KIF5B for Na_v_1.8, or local translation within the axon, as shown for Na_v_1.6 (Bao, 2015; González et al., 2016). In the CNS, nodal protein transport and targeting is poorly understood to date. Anterograde transport of Na_v_1.2 and potassium channels Kv3.1b has been linked to KIF5B, through a direct interaction with AnkG (Barry et al., 2014; Xu et al., 2014). Interaction of APP, which regulates Na_v_1.6 membrane expression (Liu et al., 2015), with kinesin-1 and dynactin-1 has been previously reported (Fu and Holzbaur, 2013).

Our data show that both kinesin-1 family and dynein/dynactin-1 axonal motors play a major role in node-like CAM axonal transport and early clusters assembly. Regarding kinesins, we demonstrated a major role of KIF5A and KIF5C in node-like CAM anterograde transport, and we hypothesize that residual activity of KIF5A in the double knockdown condition (as loss of expression was incomplete), or redundancy with KIF5B (or another kinesin motor) activity might account for residual anterograde transport after this double inactivation. As for KIF5B, although it has been reported to play a role in Na_v_1.2 transport (Barry et al., 2014), technical limitations preventing the possibility of an efficient KIF5B knockdown and a triple knockdown for all kinesin-1 motors did not allow us to determine its specific contribution in node-like CAM axonal transport.

In addition to its impact on anterograde transport, lack of KIF5A and C also resulted in decreased retrograde transport of node-like CAMs. This unexpected effect could be an adaptive response to the reduced amount of proteins in the axon. Alternatively, it could be related to modulation by molecular partners binding to both kinesin-1 and dynactin-1, such as JIP1 in the case of APP axonal transport, which plays the role of an adaptor for both kinesins and dyneins and thus regulates their transport (for review, Fu and Holzbaur, 2013; Hancock, 2014; Hendricks et al., 2010; Müller et al., 2008), or it could mediated by co-dependance mechanisms, as shown for Prp trafficking, in which dynein motility is increased by the presence of KIF5C at the vesicle (Encalada et al., 2011). The reduced number of puncta in the axon after dynactin-1 silencing could be accounted for either by delayed neuronal maturation (for review, Moughamian and Holzbaur, 2018) or by direct impact on anterograde axonal trafficking (Moughamian and Holzbaur, 2012). Such a dynactin silencing-induced delay in neuronal maturation might also partially account for the observed reduced node-like early clustering.

### *β2Na*v plays a major role in the initial step of node-like clustering

After being transported, some node-like proteins become enriched at sites located distally to existing early clusters, where new clusters form. By analyzing the chronology of these enrichments, we show that β2Na_v_ is enriched prior to other CAMs at these discrete sites. This precocious enrichment of β2Na_v_ is reminiscent of the results obtained in retinal ganglion cell cultures and P14 optic nerve sections by Kaplan et al. (Kaplan et al., 2001) showing β2Na_v_ expression in developing nodes, whereas β1Na_v_ was not expressed. Moreover, we show that miRNA mediated knock-down of β2Na_v_ decreased not only the number of GABAergic neurons forming nodal clusters by 50%, but also the number of clusters along the axons still presenting clusters (data not shown). Conversely, the overexpression of β2Na_v_ increases by 1.5-fold the number of GABAergic neurons with early clusters, which demonstrates that β2Na_v_ indeed plays a major role in node-like cluster assembly. Of note, an incomplete β2Na_v_ suppression or a compensatory increase of β1 or β4Na_v_ might account for the incomplete impact of β2Na_v_ silencing on early clustering. It is furthermore highly likely that β2Na_v_ is not the unique intrinsic organizer of early node-like clustering. In this respect, we have shown that β2Na_v_ is always associated with αNa_v_ and AnkG at these enrichment sites, and we previously reported that AnkG suppression impairs early clustering (Freeman et al., 2015). Early clustering thus appears to be initiated as the assembly of an axonal tripartite complex in the CNS rather than of a single axonal partner, as observed in the PNS for Nfasc186 (Eshed et al., 2005; Zhang et al., 2012).

β2Na_v_ function in early clustering may rely on different mechanisms. It could depend on its role in membrane targeting and stabilization of the initial node-like complex, mediated by β2Na_v_ extracellular adhesive properties and its direct interaction with αNa_v_ and AnkG (for review, Malhotra et al., 2000; Winters and Isom, 2016). Alternatively, β2Na_v_ subunit can play a role through regulation of αNa_v_ expression, and in particular Na_v_1.1, as previously described (Kim et al., 2011, 2007, 2005), this αNa_v_ isoform being expressed in GABAergic neurons forming early clusters (Freeman et al., 2015 and Desmazières, unpublished data). β2Na_v_ is furthermore sufficient to increase early clustering amongst hippocampal GABAergic neurons.

Taken together, our data converge toward a two-step mechanism for early cluster formation, with an initial loose clustering of a complex comprised of αNa_v_, β2Na_v_ and AnkG, followed by secondary recruitment of other nodal CAMs, associated with a restriction and stabilization of the structure (Fig. 7). Furthermore, recently generated data suggest that Nfasc186 could be partially recruited to forming clusters by relocation from the membrane pool (data not shown). It is possible that extracellular partners, some of them being secreted by oligodendrocytes, interact with nodal CAMs in this second step, as it has been shown that oligodendrocyte secreted factor(s) are required for node-like clusters assembly (Dubessy et al., 2019; Freeman et al., 2015; Kaplan et al., 2001, 1997; Susuki et al., 2013).

**Figure 7.**
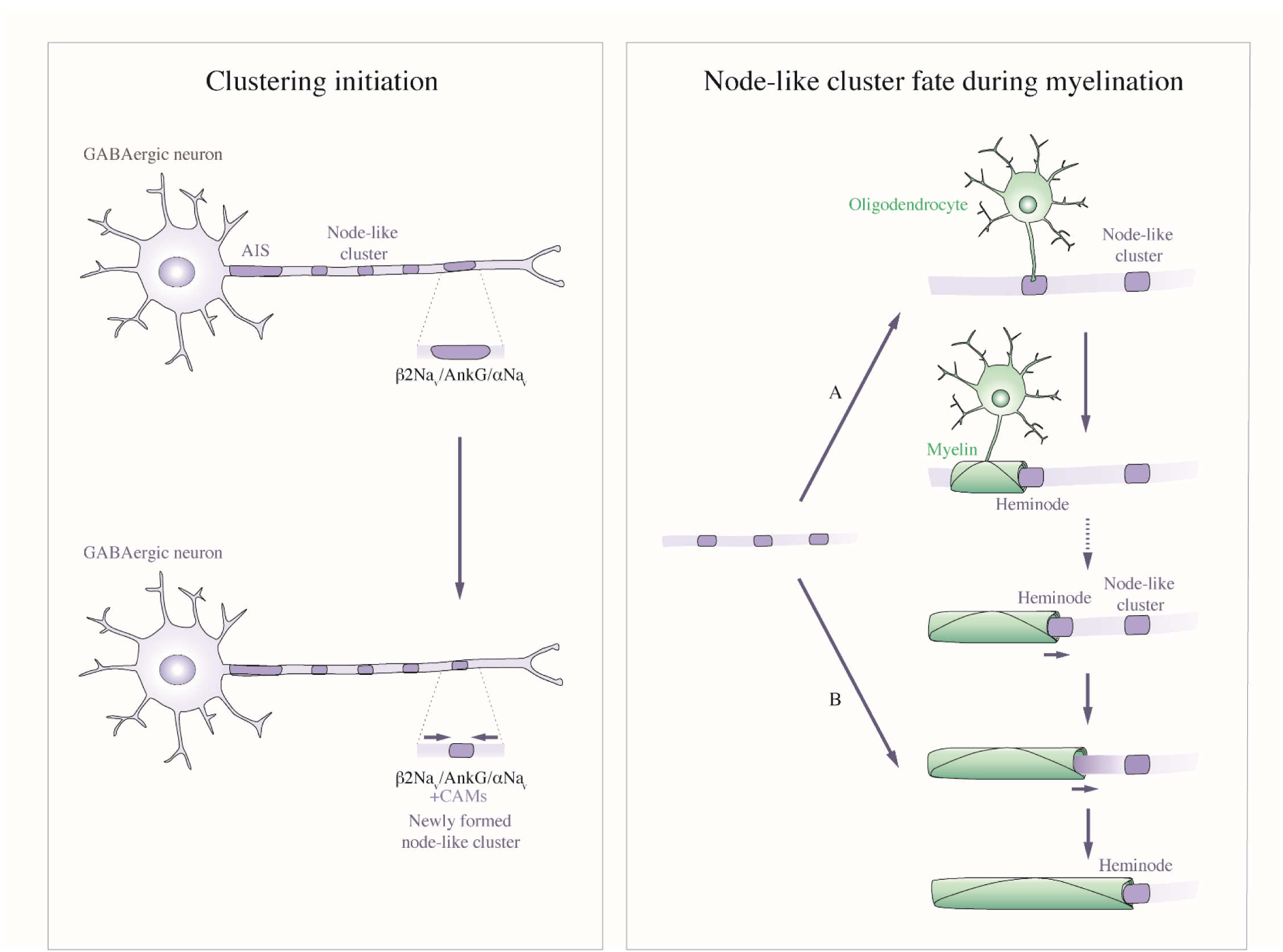
Model of node-like clustering initiation and fate during myelination. **(Left)** Prior to myelination, a tripartite complex composed of β_2_Na_v_, αNa_v_ and AnkG accumulates distally to existing node-like clusters. Recruitment of other nodal CAMs to these enrichments are associated to their restriction and stabilization. **(Right)** At the onset of myelination, node-like clusters can be contacted by oligodendrocyte processes and act as localization signal to guide myelin initiation site, thus becoming heminodal structures **(A)**. When myelination proceeds, node-like structures participate in nodal assembly by fusion with heminodes **(B)**.

### Early clusters participate in node of Ranvier formation

Having gained insight into the mechanisms underlying early cluster formation, we then addressed the fate of the node-like clusters by answering the following questions: Do early clusters persist or disappear when myelination proceeds? Also, to what extent do they participate in nodes of Ranvier formation? The mean distance between node-like clusters is about half of the distance between two mature nodes (mean distance between node-like clusters: 13.8 ± 2.1 µm by live-imaging, n=23 and 15.8 ± 0.7 µm in fixed cultures, n>400; mean distance between mature node of Ranvier: 32.7 ± 3.1 µm in fixed cultures; n>400, Freeman et al., 2015 for fixed cultures). This suggests that a reorganization of these structures must occur, with removal or fusion of some of the clusters to form mature nodes.

To further address this question, we used *in vitro* live imaging of myelinating hippocampal cultures and were able to show that node-like clusters are not removed when myelination proceeds, but persist and participate in nodal assembly. We observed multiple cases of early clusters fusing with a heminodal structure at the tip of a moving myelin sheath (Fig. 7). Interestingly, we observed a drastic reduction in myelin sheath extension velocity when the myelin tip approaches a cluster area. The partial diffusion of nodal markers (from the heminode or early cluster) prior to the fusion, together with the long “pausing time” following myelin tip apposition to the node-like cluster, suggests that axonal cytoskeletal rearrangement occurs when the myelin sheath reaches the vicinity of node-like clusters. These results are in line with the recent hypothesis of a dynamic cytoskeletal interface at the tip of the myelin, with a prominent role of the axonal cytoskeleton in oligodendrocyte process extension (Ghosh et al., 2018). Due to the long pausing time following myelin tip apposition to the early cluster, we were unable to cover the subsequent period corresponding to heminodal-heminodal fusion when myelin progresses, which has already been demonstrated as corresponding to the latest stage of nodes of Ranvier formation (Brivio et al., 2017; Ghosh et al., 2018; Zonta et al., 2008).

It has been shown that formation of myelin segment of adequate length can occur along synthetic fibers of appropriate diameter *in vitro*, suggesting no axonal signal is required for myelin initiation and elongation in this context (Bechler et al., 2015). The striking detection of the onset of myelin deposition at the edge of early node-like clusters prompted us to hypothesize an unexpected role for these clusters in specifying the site for myelin initiation (Fig. 7). This hypothesis was recently reinforced by preliminary data generated using an *ex vivo* live imaging approach on cerebellar slices, in which early clusters form along Purkinje cells axons prior to myelin formation and where we observed the initiation of myelin deposition at the node-like cluster vicinity. This suggests the existence of underlying axonal signals of attraction/repulsion of oligodendrocytes processes at the node-like cluster which could lead to a preferential initiation site for myelin deposition. Whether they might be linked to neuronal electrical activity or to locally secreted factors is still to be characterized. The fact that early node-like clustering also depends on oligodendroglial secreted cues (Dubessy et al., 2019; Freeman et al., 2015; Kaplan et al., 2001, 1997) further shows that a tightly regulated dual interaction between neuron and glia is required for myelinated fiber organization.

## MATERIALS AND METHODS

The care and use of animals in all experiments conform to institutional policies and guidelines (UPMC, INSERM, French and European Community Council Directive 86/609/EEC).

### Animals

The following rat and mouse strains were used in this study: Sprague–Dawley or Wistar rats (Janvier Breeding Center, Saint-Berthevi, France) and PLP-GFP mice (Spassky et al., 2001).

### Cell culture

Mixed hippocampal cultures (containing neurons and glial cells) were prepared according to procedures described in (Freeman et al., 2015). Briefly, pooled hippocampi of E18.5 rat embryos were dissociated enzymatically by trypsin treatment (0.1%; Worthington) for 20 min with DNase (50 μg/mL, Worthington). After trypsin neutralization, cells were centrifuged at 1500 rpm for 5 min, resuspended, and then seeded on polyethylenimine precoated glass coverslips or on 35 mm glass-bottom dishes (81158; Ibidi, BioValley) at a density of 5.0 × 10^4^ cells/35 mm^2^. Cultures were maintained for 24 h in a 1:1 mixture of DMEM-Glutamax (31966047; Gibco, Thermo Fisher Scientific) with 10% FCS (100 IU/mL), penicillin-streptomycin (100 IU/mL each), and neuron culture medium (NCM). Culture medium was then replaced by a 1:1 mixture of Bottenstein–Sato (BS) medium with PDGF-A (0.5%) and NCM, and subsequently, one-half of the medium was renewed every 3 days and replaced by NCM.

For hippocampal neuron-oligodendrocyte myelinating co-culture, 5.0 × 10^4^ cells FACS-sorted oligodendrocytes from PLP-GFP mice (Spassky et al., 2001) were added at 14 DIV per well, and half of the medium was replaced with differentiation medium to induce myelination. From this point onward, we used differentiation medium without phenol red for live-imaging.

### Media composition

#### NCM

Neurobasal Medium (21103–049; Gibco, Thermo Fisher Scientific) supplemented with 0.5 mM L-glutamine, B27 (1X; Invitrogen, Thermo Fisher Scientific), and penicillin-streptomycin (100 IU/mL each).

#### BS medium

DMEM-Glutamax supplemented with transferrin (50 μg/mL), albumin (50 μg/mL), insulin (5 μg/mL), progesterone (20 nM), putrescine (16 μg/mL), sodium selenite (5 ng/mL), T3 (40 ng/mL), and T4 (40 ng/mL).

#### Differentiation medium

DMEM-Glutamax supplemented with transferrin (50 μg/mL), albumin (50 μg/mL), insulin (5 μg/mL), putrescine (16 μg/mL), sodium selenite (5 ng/mL), T3 (40 ng/mL), biotin (10 ng/mL), vitamin B12 (27.2 ng/mL), ceruloplasmin (100 ng/mL), hydrocortisone (0.05 μM), CNTF (0.1 ng/mL), and sodium pyruvate (1 mM).

### Plasmid Constructs

In order to express nodal proteins tagged with fluorescent markers, the coding sequence of the nodal markers placed in frame with the mCherry or GFP coding sequence at their 3’ end (corresponding to C-ter in the protein) was inserted into the pEntr vector. The sequence of interest was then recombined into the pTRIP-Syn (human synapsin promoter inserted into the pDEST-Rfa-DeltaU3, gift from Dr. P. Ravassard, ICM, France, ICM, France) using Gateway LR clonase II (Thermo Fischer Scientific), allowing the fluorescently-tagged nodal protein expression specifically in neurons. The pCMVmCherry-N1 and pEGFP-N1 were purchased from Clontech. The coding sequences for β1Na_v_-GFP and β2Na_v_-YFP were also purchased commercially (Origene, MG202467, and Biovalley, DQ895640, respectively), the pCMVNfasc186-mCherry and the pCMVAPP-mCherry were kindly provided by Pr. P.J. Brophy, University of Edinburgh, UK, and Dr. M-C. Potier, ICM, France, respectively. pEntrmCherry was generated using KpnI/XhoI digest to retrieve mCherry from pCMVmCherry-N1 and insert it into pEntr. pEntrNfasc186-mCherry was generated by EcoRI/XhoI digestion of pEntr and insertion of Nfasc186-mCherry retrieved from pCMVNfasc186-mCherry by EcoRI/SalI digestion. β1Na_v_-mCherry and β2Na_v_-mCherry coding sequences were generated by cloning a PCR fragment encoding β1Na_v_ or β2Na_v_ without stop codon (for primers see Table S1) into pEntrNfasc186-mCherry after removal of the Nfasc186 coding sequence using EcoRI and HindIII sites. pEntrβ2Na_v_-GFP was generated by digestion of pEntrβ2Na-vmCherry with HindIII/KpnI to retrieve mCherry and insertion of GFP following PCR expansion using pEGFP-N1 as matrix (for primers see Table S1). The pTRIP plasmids generated were further used to generate lentiviral vectors (ICM vectorology platform). The miRNA sequences are indicated in Table S1. The miRNA plasmids used for knockdown studies were generated following manufacturer’s instructions (Thermo Fisher Scientific, K4936-00). The miRNA SCN2b plasmid has been provided by Dr. N. Sol-Foulon.

### Transfection and Lentiviral Infection

Transfection of rat primary neurons was performed at either 5 or 6 days *in vitro* (DIV) with a total of 500 ng DNA and 1.0 μL Lipofectamine 2000 reagent per well (Thermo Fisher Scientific) in Opti-MEM reduced serum medium (Thermo Fisher Scientific). 50 to 100 ng/μL of plasmid were used for nodal protein expression and 400 ng/μL for miRNA (supplemented with pBlueScript). For viral transduction of neurons, lentiviruses were added for 2 hrs to cultures at 10^5^ VP/μl at either 5 or 6 DIV. Transfected or transduced neurons were fixed at 14, 17 or 21 DIV for immunocytochemical analysis. Live cultures were used at either 10 or 17 DIV for axonal transport studies and from 21 DIV to 24 DIV for node-like cluster fate studies.

### Immunofluorescence

Cell cultures were fixed with 4% PFA for 10 min, or with 1% PFA for 10 min at room temperature (RT) and then incubated with methanol for 10 min at −20 °C (for αNa_v_ staining). After fixation, cells were washed in PBS, then incubated with blocking solution (PBS with 5% Normal Goat Serum (50-062Z; Thermo Fisher Scientific), 0.1% Triton X-100) for 15 min and with primary antibodies solutions for 2 hrs at RT. Coverslips were then washed in PBS and incubated with secondary antibodies solutions for 1 hr at RT. Coverslips were washed in PBS and mounted on glass slides with Fluoromount G (Southern Biotech).

### Antibodies

The following mouse primary antibodies were used: mouse IGg2a anti-AnkyrinG (clone N106/36; 1:100), Nfasc (pan, external; clone A12/18; 1:100) from Antibodies Incorporated (Neuromab); mouse IgG1 anti-Na_v_ (pan; clone K58/35; 1:250) from Sigma; mouse IgG2a anti-GAD67 (clone 1610.2; 1:400) and mouse IGg1 anti-MAP2 (1:1000) from Millipore. Rabbit polyclonal primary antibodies included anti-β1Na_v_ (1:100; kindly provided by Pr. P.J. Brophy, University of Edinburgh, UK), β2Na_v_ (1:250; Millipore), Nfasc (pan; 1:100; Abcam), AnkG (1:500; kindly provided by Pr. F. Couraud, INSERM, UMRS952, Paris), kinesin 5A (1:500; Abcam), kinesin 5B (1:500; Abcam), kinesin 5C (1:500; Abcam), KIF3A (1:100, Abcam), KIF4A (1:100, Abcam), dynactin-1 (1:1000; abcam); Neurofilament M (1:400; Millipore), Caspr (1:500; Abcam). We also used chicken anti-GFP (1:400; Millipore); m-cherry (1:2000, EnCor) and rat anti-PLP (1:10; hybridoma; kind gift from du Dr. K. Ikenaka, Okasaki, Japan). Secondary antibodies were goat anti-rabbit, anti-mouse IgG2a, IgG1, anti-chicken, or anti-rat coupled to Alexa Fluor 488, 594 or 647 from Invitrogen (1:1000).

### Fluorescent-activated cell sorting purification of GFP-positive oligodendrocytes

Isolation was performed in two steps, as described previously (Piaton et al., 2011). Briefly, brains from P7-P10 *PLP-GFP* mice pups (Spassky et al., 2001) were used to obtain oligodendrocytes. Tissue was dissected in 1X HBSS supplemented with 0.01 mM HEPES buffer, 0.75% sodium bicarbonate (Invitrogen), and 1% penicillin/streptomycin and mechanically dissociated. Following a second step corresponding to enzymatic dissociation using papain (30 μg/ml in DMEM-Glutamax, with 0.24 μg/ml l-cystein and 40 μg/ml DNase I), cells were put on a preformed Percoll density gradient before centrifugation for 15 min. Cells were then collected and stained with propidium iodide (PI) for 2 min at room temperature (Moyon et al., 2015). In a second step, GFP-positive and PI-negative cells were sorted by fluorescence-activated cell sorting (FACS; Aria, Becton Dickinson) and collected in pure fetal bovine serum. For cultures, cells were resuspended in differentiation medium before being added on top of mixed hippocampal cells cultured in glass bottom dishes (81158, Ibidi).

### Western Blot

PC12 cells or N2a cells were cultured in DMEM-Glutamax (31966047; Gibco, Thermo Fisher Scientific) with 10% FCS (100 IU/mL) and penicillin-streptomycin (100 IU/mL each). Cells were transfected at 70% confluence using Lipofectamine 2000 reagent (Thermo Fisher Scientific) according to manufacturer’s instructions and harvested 48 hrs after transfection. Cells were washed once in 1X PBS on ice and lysed in 1X RIPA buffer (Sigma), supplemented with protease inhibitors (Roche). Cell lysates were centrifuged at 20.000 rpm for 20 mins and the supernatants were collected. Protein concentrations were determined by the BCA method (Thermo Fisher Scientific) and protein samples (30-50 μg) were separated on 4%–12% Bis-Tris gradient gels (Thermo Fisher Scientific) and transferred to nitrocellulose membranes (Whatman). Non-specific antigen binding was blocked with 10% milk in TBS-T (50 mM Tris, 150 mM NaCl and 0.1% Tween-20) for 1 hr.

Membranes were then incubated with primary antibodies (anti-kinesin 5A (1:500; Abcam), anti-kinesin 5B (1:500; Abcam), anti-KIF4A (1:75, Abcam) or anti-β-Tubulin (1:1000, Abcam)) in blocking solution overnight at 4°C. After washing, the blots were incubated with horseradish peroxidase–conjugated anti-mouse, anti-rabbit or anti-goat secondary antibodies (all used at 1:1000) for 2 hrs, followed by washes in TBS-T. Immunoblots were developed with SuperSignal chemiluminescent substrate (GE Healthcare), imaged using the ChemiDoc MP Imaging System (Bio-Rad) and analyzed with ImageLab software (Bio-Rad).

### Imaging

#### Confocal microscopy

Confocal microscopy was performed using an Olympus FV-1000 or FV-1200 Upright Confocal Microscope with a 1.35 numerical aperture 63x objective (oil immersion). We used 473, 545 and 635-nm laser wavelength outputs with laser intensities of 17 mW, 5 mW and 23 mW, respectively. Z-series of optical sections were performed at 0.3 μm increments. Green, red and far-red fluorescence were acquired sequentially. Maximum intensity orthogonal projection of images and adjustment for brightness and contrast were carried out using ImageJ software (NIH, Bethesda, Maryland).

#### Videomicroscopy

For live imaging, cells were grown on 35 mm glass-bottom dishes (81158; Ibidi, BioValley) and transfected as described previously. For live imaging, dishes were placed in a temperature-controlled imaging chamber stabilized at 37° under 5% CO2. Single and dual live-imaging were performed on an AxioVert200 inverted Zeiss microscope using a 63x PlanApo oil objective (numerical aperture = 1.4), and captured using a Photometrics DualCam2 Evolve camera through GFP, mCherry or GFP/mCherry filter sets. We used a mercury lamp (Osram HBO 100W/2). Images were taken 30 to 50 μm distally to the axon initial segment (AIS) in streaming mode with a 250 ms exposure time for 201 frames using Metamorph software (Molecular Devices). Specific parameters (e.g. time between frames, duration of imaging) are indicated in the figure legends. Axons were identified morphologically by the lack of dendritic spines and the enrichment of the fluorescently-tagged protein at the AIS. To prevent phototoxicity during live-imaging, we used culture-medium without phenol red and low lamp intensity. Media pH was monitored before and after imaging.

#### Imaging of node-like cluster fate

To detect nodal structures in myelinating co-cultures, cells were incubated for 1 hr prior to imaging, with an antibody targeting an external epitope of Neurofascin (1:1000, pan, external; clone A12/18; NeuroMab) directly coupled to an Alexa Fluor 594 (A10474; Thermo Fisher Scientific). Imaging was carried out on a Yokogawa CSU-X1 M Spinning Disk, mounted on an inverted Leica DMi8 microscope, with a 40x 1.30 oil immersion objective, using the 488 nm and 561 nm laser lines (with laser intensities of 660 μW and 70 μW, respectively) and a Hamamatsu Flash 4 LT camera. Images were acquired at 37°C in a temperature-controlled chamber under 5% CO2 using Metamorph software (Molecular Devices), for periods of 5 to 54 hrs with confocal Z-series stack acquired for both GFP and AlexaFluor594 at 150 ms exposure, and *z*-intervals of 0.5 μm between optical slices. Specific parameters (e.g. time between frames, duration of imaging) are indicated in the figure legends. To prevent phototoxicity during live-imaging, we used culture-medium without phenol red and low lamp intensity. Media pH was monitored before and after imaging.

To address whether a node-like structure is removed when myelin progresses towards it or whether the node like structure would be integrated to the heminode at the tip of the myelin, we focused on area of the culture where we could observe a node like cluster in the vicinity of a myelinating sheath associated to a heminode (at a distance of about 10 μm of the node-like cluster) and live imaged this area up to 50 hrs.

### Analysis

#### Cell Image Analysis

For the analysis of single or dual-color live imaging, kymographs (representing motion of fluorescent structures over time) were generated with the ImageJ software using the KymoToolBox plugin (kindly provided by Dr. F. Cordelière, University of Bordeaux). For analysis, individual trajectories were manually traced. For dual-color imaging, trajectories were traced for each of the GFP and mCherry kymographs, binarized, pseudocolored (green and red respectively) and merged. For all kymographs, moving puncta [anterograde (toward the axon terminal), retrograde (toward the cell body), or bidirectional] were defined as having instantaneous speeds above 0.1 μm/s and covering a total distance of more than 3 μm in a given direction. The mean velocity was defined as the mean of the instantaneous velocities calculated for each fragment of a trajectory as the Δdistance covered/Δtime. We also calculated the mean number of puncta per 100 μm in each condition and the distribution of anterograde, retrograde, bidirectional and stationary puncta. Group analysis of the trajectories was performed using a homemade Excel macro by Dr. J. Tailleur (University of Paris Diderot).

#### Quantification of neurons with clusters

At least 50 neurons per coverslip, identified by the presence of an AIS, were counted, and the percentage of neurons with αNa_v_/AnkG clusters was determined for at least 3 coverslips per condition. Results were expressed as means ± SEMs of at least three independent experiments. αNa_v_/AnkG clusters were defined as described in Freeman, et al., 2015 (Freeman et al., 2015) (i.e. size of clusters less than 8 μm).

#### Time-lapse analysis

Following live imaging, the cultures were fixed and stained with PLP, to confirm that imaged nodal structures were indeed node-like clusters, i.e that these clusters of nodal proteins were not flanked by GFP-negative myelin (endogenous to the culture, not shown). These experiments were repeated using lentiviral transduction to express Nfasc186-mCherry, to exclude a potential bias due to the use of the Nfasc186 external epitope-directed antibody.

For time-lapse acquisitions, using ImageJ software, we performed maximum-intensity projection of image stacks and alignment between time points (using StackReg Plugin). Median filter and subtraction of background were applied to remove noise and images were adjusted for brightness and contrast. Myelin progression measurements were made in two dimensions from maximum-intensity projections of image stacks along time.

#### Measurements of myelin sheath growth

For the analysis of myelin sheath growth over time, we defined t0 as the first time point acquired during an imaging session. The final time was defined as the end of the imaging for myelin sheaths without node like clusters at their vicinity or the time point when a heminode and a node-like cluster fused (as no further myelin growth was observed lateron). To assess the distance covered by a myelin sheath over time, we measured the distance between the tip of myelin sheath and the center of the facing node like cluster at each time point. In the absence of a node-like cluster, we chose a morphological landmark along the axon as a reference to measure myelin growth. Data are expressed as the distances covered between two time-points cumulated along time.

#### Statistical analysis

Statistical analysis was performed using either *R* or Prism (GraphPad) software.

For mean number of puncta per 100 μm (Fig. 1, 2, 3; Fig. S1, S3) and mean speed analysis (Fig. S1), groups were compared by unpaired *t*–test. For the grouped analysis, multiple groups were compared by one-way analysis of variance (ANOVA) test, with a Tukey’s post-test. For miRNA studies (Fig. 2, 3, 4, 5, S3 and S4) significance was assessed by unpaired *t*–test. For the distribution analysis (Fig. 1 D, 2, 3 and Fig. S1 D and S3 C-E and S3 I), the difference between proteins across all types of movement was assessed using a Chi-squared test. Statistical analysis of myelin progression measurements was performed using a Mann-Whitney non-parametric *t*-test. *p-*values, including for grouped analysis, are indicated in the Table S2.

## ACKNOWLEDGMENTS

We thank Dr. M. Gomes-Pereira, Dr. M.-C. Potier and Dr. P. Ravassard for their gift of the pCMV-Synaptophysin-DsRed, pCMV-APP-mCherry and pTrip-Syn and pCMV-GFP plasmids respectively. We thank Dr. B. Zalc, Dr. N. Sol-Foulon and Dr. M. Davenne for insights and discussion. We thank the icm.Quant imaging platform, Dr. D. Langui and D. Akbar for their support in videomicroscopy acquisitions. We thank the ICM biostatistics platform (iCONICS) for support in statistical analysis. We thank the CELIS, vectorology, genotyping and PhenoICMouse ICM facilities. This work was funded by INSERM, ICM, ARSEP Grant R13123DD (to C.L. and A.D.), FRM fellowship, SPF20110421435 (to A.D.), FDT20170437332 (to M.T.), Prix Bouvet-Labruyère - Fondation de France (to A.D.), ANR JC (ANR-17-CE16-0005-01; to A.D), NRJ foundation and FRC (« Espoir en tête », Rotary Club).

## AUTHOR CONTRIBUTION

M. Thetiot, S.A. Freeman, T. Roux, C. Lubetzki, A. Desmazières designed research; M. Thetiot, S.A. Freeman, T. Roux., A.L. Dubessy, M.S. Aigrot., Q. Rappeneau, A. Desmazières performed research and analyzed data; J. Tailleur, N. Sol-Foulon provided tools and inputs; M. Thetiot and F.X. Lejeune performed biostatistical analysis. M. Thetiot, S.A. Freeman, C. Lubetzki, and A. Desmazières wrote the paper.

The authors declare no conflict of interest.

## TABLE SUPPLEMENT LEGENDS

**Table S1.** Oligonucleotides used for constructs generation.

**Table S2.** Biostatistical analyses.

## MOVIE LEGENDS

**Movie S1. Nodal proteins co-transport *in vitro* (related to Figure 1).**

Video shows the dynamic movement of β2Na_v_-GFP (first panel) and β1Na_v_-mCherry (second panel) in axons of 17 DIV rat hippocampal neurons. For each panel, rows showing color dots correspond to the motion of tracked vesicles. Green dots are for β2Na_v_-GFP vesicles movements, red are for β1Na_v_-mCherry vesicles movements, and yellow are for co-transported vesicles (bottom panel). Retrograde motility is toward the left and anterograde motility is toward the right. Time-lapse images were acquired at a rate of 250 ms per frame for 50 s using a videomicroscope (AxioVert200; Carl Zeiss). Videos are displayed at a rate of 10 frames/s. Time stamp indicates minutes and seconds. Scale bar, 5 µm.

**Movie S2. *β2Na*v accumulates prior to the other CAMs (related to Figure 5).**

Video shows two local accumulations of a subset of β2Na_v_-GFP puncta (top panel) while β1Na_v_-mCherry vesicles exhibit dynamic movement (bottom panel). Time-lapse images were acquired at a rate of 250 ms per frame for 50 s using a videomicroscope (AxioVert200; Carl Zeiss). Videos are displayed at a rate of 10 frames/s. Time stamp indicates minutes and seconds. Scale bar, 5 µm.

**Movie S3. Initiation of early clustering.**

Movie showing two axonal enrichments of Nfasc186 (red) and their restriction along time to form stable early clusters. Cultures were imaged at 20 DIV every hour for 34 hrs. Video is displayed at a rate of 3 frames/s. Scale bar, 5 µm.

**Movie S4. Myelin apposition at an early cluster (related to Figure 6).**

Time-lapse movie showing an early structure (Nfasc186, red) being contacted by an oligodendrocyte process followed by myelin deposition (green). Cultures were imaged at 21 DIV every hour for 3 hrs. Video is displayed at a rate of 2.5 frames/s. Scale bar, 5 µm.

**Movie S5. Early structures guide myelin deposition (related to Figure 6).**

Time-lapse movie showing an oligodendrocyte process (green) myelinating at the close vicinity of an early cluster (Nfasc186, red). Cultures were imaged at 21 DIV every hour for 30 hrs. Video is displayed at a rate of 5 frames/s. Scale bar, 5 µm.

**Movie S6. Heminode fusion with early cluster during myelination (related to Figure 3).**

Time-lapse movie showing the myelin tip (green) flanked with a heminode (Nfasc186, red) progressing along the axon and reaching an early cluster (Nfasc186, red). Cultures were imaged at 21 DIV every hour for 43 hrs. Video is displayed at a rate of 7 frames/s. Scale bar, 5 µm.

**Movie S7. Heminode fusion with an early cluster during myelination (related to Figure 6).**

Time-lapse movie showing myelin progression (green) with a partial diffusion of the heminode (Nfasc186, red) before fusing with an early cluster (Nfasc186, red). Cultures were imaged at 21 DIV every hour for 23 hrs. Video is displayed at a rate of 3 frames/s. Scale bar, 5 µm.

## REFERENCES

Amor, V., Zhang, C., Vainshtein, A., Zhang, A., Zollinger, D.R., Eshed-Eisenbach, Y., Brophy, P.J., Rasband, M.N., Peles, E., 2017. The paranodal cytoskeleton clusters Na+ channels at nodes of Ranvier. eLife 6. https://doi.org/10.7554/eLife.21392

Bao, L., 2015. Trafficking regulates the subcellular distribution of voltage-gated sodium channels in primary sensory neurons. Mol. Pain 11, 61. https://doi.org/10.1186/s12990-015-0065-7

Barry, J., Gu, Y., Jukkola, P., O’Neill, B., Gu, H., Mohler, P.J., Rajamani, K.T., Gu, C., 2014. Ankyrin-G directly binds to kinesin-1 to transport voltage-gated Na+ channels into axons. Dev. Cell 28, 117–131. https://doi.org/10.1016/j.devcel.2013.11.023

Bechler, M.E., Byrne, L., ffrench-Constant, C., 2015. CNS Myelin Sheath Lengths Are an Intrinsic Property of Oligodendrocytes. Curr. Biol. 25, 2411–2416. https://doi.org/10.1016/j.cub.2015.07.056

Bonetto, G., Hivert, B., Goutebroze, L., Karagogeos, D., Crépel, V., Faivre-Sarrailh, C., 2019. Selective Axonal Expression of the Kv1 Channel Complex in Pre-myelinated GABAergic Hippocampal Neurons. Front. Cell. Neurosci. 13. https://doi.org/10.3389/fncel.2019.00222

Boyle, M.E., Berglund, E.O., Murai, K.K., Weber, L., Peles, E., Ranscht, B., 2001. Contactin orchestrates assembly of the septate-like junctions at the paranode in myelinated peripheral nerve. Neuron 30, 385–397.

Bréchet, A., Fache, M.-P., Brachet, A., Ferracci, G., Baude, A., Irondelle, M., Pereira, S., Leterrier, C., Dargent, B., 2008. Protein kinase CK2 contributes to the organization of sodium channels in axonal membranes by regulating their interactions with ankyrin G. J. Cell Biol. 183, 1101–1114. https://doi.org/10.1083/jcb.200805169

Brivio, V., Faivre-Sarrailh, C., Peles, E., Sherman, D.L., Brophy, P.J., 2017. Assembly of CNS Nodes of Ranvier in Myelinated Nerves Is Promoted by the Axon Cytoskeleton. Curr. Biol. CB 27, 1068–1073. https://doi.org/10.1016/j.cub.2017.01.025

Buttermore, E.D., Thaxton, C.L., Bhat, M.A., 2013. Organization and maintenance of molecular domains in myelinated axons. J. Neurosci. Res. 91, 603–622. https://doi.org/10.1002/jnr.23197

Chen, C., Bharucha, V., Chen, Y., Westenbroek, R.E., Brown, A., Malhotra, J.D., Jones, D., Avery, C., Gillespie, P.J., Kazen-Gillespie, K.A., Kazarinova-Noyes, K., Shrager, P., Saunders, T.L., Macdonald, R.L., Ransom, B.R., Scheuer, T., Catterall, W.A., Isom, L.L., 2002. Reduced sodium channel density, altered voltage dependence of inactivation, and increased susceptibility to seizures in mice lacking sodium channel beta 2-subunits. Proc. Natl. Acad. Sci. U. S. A. 99, 17072–17077. https://doi.org/10.1073/pnas.212638099

Çolakoğlu, G., Bergstrom-Tyrberg, U., Berglund, E.O., Ranscht, B., 2014. Contactin-1 regulates myelination and nodal/paranodal domain organization in the central nervous system. Proc. Natl. Acad. Sci. U. S. A. 111, E394–403. https://doi.org/10.1073/pnas.1313769110

Coman, I., Aigrot, M.S., Seilhean, D., Reynolds, R., Girault, J.A., Zalc, B., Lubetzki, C., 2006. Nodal, paranodal and juxtaparanodal axonal proteins during demyelination and remyelination in multiple sclerosis. Brain J. Neurol. 129, 3186–3195. https://doi.org/10.1093/brain/awl144

Craner, M.J., Newcombe, J., Black, J.A., Hartle, C., Cuzner, M.L., Waxman, S.G., 2004. Molecular changes in neurons in multiple sclerosis: altered axonal expression of Nav1.2 and Nav1.6 sodium channels and Na+/Ca2+ exchanger. Proc. Natl. Acad. Sci. U. S. A. 101, 8168–8173. https://doi.org/10.1073/pnas.0402765101

Davis, J.Q., Lambert, S., Bennett, V., 1996. Molecular composition of the node of Ranvier: identification of ankyrin-binding cell adhesion molecules neurofascin (mucin+/third FNIII domain-) and NrCAM at nodal axon segments. J. Cell Biol. 135, 1355–1367.

Desmazieres, A., Zonta, B., Zhang, A., Wu, L.-M.N., Sherman, D.L., Brophy, P.J., 2014. Differential stability of PNS and CNS nodal complexes when neuronal neurofascin is lost. J. Neurosci. Off. J. Soc. Neurosci. 34, 5083–5088. https://doi.org/10.1523/JNEUROSCI.4662-13.2014

Dubessy, A.-L., Mazuir, E., Rappeneau, Q., Ou, S., Abi Ghanem, C., Piquand, K., Aigrot, M.-S., Thétiot, M., Desmazières, A., Chan, E., Fitzgibbon, M., Fleming, M., Krauss, R., Zalc, B., Ranscht, B., Lubetzki, C., Sol-Foulon, N., 2019. Role of a Contactin multi-molecular complex secreted by oligodendrocytes in nodal protein clustering in the CNS. Glia. https://doi.org/10.1002/glia.23681

Dzhashiashvili, Y., Zhang, Y., Galinska, J., Lam, I., Grumet, M., Salzer, J.L., 2007. Nodes of Ranvier and axon initial segments are ankyrin G-dependent domains that assemble by distinct mechanisms. J. Cell Biol. 177, 857–870. https://doi.org/10.1083/jcb.200612012

Encalada, S.E., Szpankowski, L., Xia, C., Goldstein, L.S.B., 2011. Stable kinesin and dynein assemblies drive the axonal transport of mammalian prion protein vesicles. Cell 144, 551–565. https://doi.org/10.1016/j.cell.2011.01.021

Eshed, Y., Feinberg, K., Carey, D.J., Peles, E., 2007. Secreted gliomedin is a perinodal matrix component of peripheral nerves. J. Cell Biol. 177, 551–562. https://doi.org/10.1083/jcb.200612139

Eshed, Y., Feinberg, K., Poliak, S., Sabanay, H., Sarig-Nadir, O., Spiegel, I., Bermingham, J.R., Peles, E., 2005. Gliomedin mediates Schwann cell-axon interaction and the molecular assembly of the nodes of Ranvier. Neuron 47, 215–229. https://doi.org/10.1016/j.neuron.2005.06.026

Fache, M.-P., Moussif, A., Fernandes, F., Giraud, P., Garrido, J.J., Dargent, B., 2004. Endocytotic elimination and domain-selective tethering constitute a potential mechanism of protein segregation at the axonal initial segment. J. Cell Biol. 166, 571–578. https://doi.org/10.1083/jcb.200312155

Feinberg, K., Eshed-Eisenbach, Y., Frechter, S., Amor, V., Salomon, D., Sabanay, H., Dupree, J.L., Grumet, M., Brophy, P.J., Shrager, P., Peles, E., 2010. A glial signal consisting of gliomedin and NrCAM clusters axonal Na+ channels during the formation of nodes of Ranvier. Neuron 65, 490–502. https://doi.org/10.1016/j.neuron.2010.02.004

Freeman, S.A., Desmazières, A., Simonnet, J., Gatta, M., Pfeiffer, F., Aigrot, M.S., Rappeneau, Q., Guerreiro, S., Michel, P.P., Yanagawa, Y., Barbin, G., Brophy, P.J., Fricker, D., Lubetzki, C., Sol-Foulon, N., 2015. Acceleration of conduction velocity linked to clustering of nodal components precedes myelination. Proc. Natl. Acad. Sci. U. S. A. 112, E321–328. https://doi.org/10.1073/pnas.1419099112

Fu, M., Holzbaur, E.L.F., 2013. JIP1 regulates the directionality of APP axonal transport by coordinating kinesin and dynein motors. J. Cell Biol. 202, 495–508. https://doi.org/10.1083/jcb.201302078

Ghosh, A., Sherman, D.L., Brophy, P.J., 2018. The Axonal Cytoskeleton and the Assembly of Nodes of Ranvier. Neurosci. Rev. J. Bringing Neurobiol. Neurol. Psychiatry 24, 104–110. https://doi.org/10.1177/1073858417710897

González, C., Cánovas, J., Fresno, J., Couve, E., Court, F.A., Couve, A., 2016. Axons provide the secretory machinery for trafficking of voltage-gated sodium channels in peripheral nerve. Proc. Natl. Acad. Sci. U. S. A. 113, 1823–1828. https://doi.org/10.1073/pnas.1514943113

Griggs, R.B., Yermakov, L.M., Susuki, K., 2017. Formation and disruption of functional domains in myelinated CNS axons. Neurosci. Res. 116, 77–87. https://doi.org/10.1016/j.neures.2016.09.010

Hancock, W.O., 2014. Bidirectional cargo transport: moving beyond tug of war. Nat. Rev. Mol. Cell Biol. 15, 615–628. https://doi.org/10.1038/nrm3853

Hendricks, A.G., Perlson, E., Ross, J.L., Schroeder, H.W., Tokito, M., Holzbaur, E.L.F., 2010. Motor coordination via a tug-of-war mechanism drives bidirectional vesicle transport. Curr. Biol. CB 20, 697–702. https://doi.org/10.1016/j.cub.2010.02.058

Howe, C.L., Mobley, W.C., 2005. Long-distance retrograde neurotrophic signaling. Curr. Opin. Neurobiol. 15, 40–48. https://doi.org/10.1016/j.conb.2005.01.010

Huxley, A.F., Stampfli, R., 1949. Evidence for saltatory conduction in peripheral myelinated nerve fibres. J. Physiol. 108, 315–339.

Jamann, N., Jordan, M., Engelhardt, M., 2018. Activity-dependent axonal plasticity in sensory systems. Neuroscience 368, 268–282. https://doi.org/10.1016/j.neuroscience.2017.07.035

Jenkins, P.M., Kim, N., Jones, S.L., Tseng, W.C., Svitkina, T.M., Yin, H.H., Bennett, V., 2015. Giant ankyrin-G: a critical innovation in vertebrate evolution of fast and integrated neuronal signaling. Proc. Natl. Acad. Sci. U. S. A. 112, 957–964. https://doi.org/10.1073/pnas.1416544112

Jenkins, S.M., Bennett, V., 2002. Developing nodes of Ranvier are defined by ankyrin-G clustering and are independent of paranodal axoglial adhesion. Proc. Natl. Acad. Sci. U. S. A. 99, 2303–2308. https://doi.org/10.1073/pnas.042601799

Kaplan, M.R., Cho, M.H., Ullian, E.M., Isom, L.L., Levinson, S.R., Barres, B.A., 2001. Differential control of clustering of the sodium channels Na(v)1.2 and Na(v)1.6 at developing CNS nodes of Ranvier. Neuron 30, 105–119.

Kaplan, M.R., Meyer-Franke, A., Lambert, S., Bennett, V., Duncan, I.D., Levinson, S.R., Barres, B.A., 1997. Induction of sodium channel clustering by oligodendrocytes. Nature 386, 724–728. https://doi.org/10.1038/386724a0

Kazen-Gillespie, K.A., Ragsdale, D.S., D’Andrea, M.R., Mattei, L.N., Rogers, K.E., Isom, L.L., 2000. Cloning, localization, and functional expression of sodium channel beta1A subunits. J. Biol. Chem. 275, 1079–1088.

Kim, D.Y., Carey, B.W., Wang, H., Ingano, L.A.M., Binshtok, A.M., Wertz, M.H., Pettingell, W.H., He, P., Lee, V.M.-Y., Woolf, C.J., Kovacs, D.M., 2007. BACE1 regulates voltage-gated sodium channels and neuronal activity. Nat. Cell Biol. 9, 755–764. https://doi.org/10.1038/ncb1602

Kim, D.Y., Gersbacher, M.T., Inquimbert, P., Kovacs, D.M., 2011. Reduced sodium channel Na(v)1.1 levels in BACE1-null mice. J. Biol. Chem. 286, 8106–8116. https://doi.org/10.1074/jbc.M110.134692

Kim, D.Y., Ingano, L.A.M., Carey, B.W., Pettingell, W.H., Kovacs, D.M., 2005. Presenilin/gamma-secretase-mediated cleavage of the voltage-gated sodium channel beta2-subunit regulates cell adhesion and migration. J. Biol. Chem. 280, 23251–23261. https://doi.org/10.1074/jbc.M412938200

Lai, H.C., Jan, L.Y., 2006. The distribution and targeting of neuronal voltage-gated ion channels. Nat. Rev. Neurosci. 7, 548–562. https://doi.org/10.1038/nrn1938

Leterrier, C., Brachet, A., Dargent, B., Vacher, H., 2011. Determinants of voltage-gated sodium channel clustering in neurons. Semin. Cell Dev. Biol. 22, 171–177. https://doi.org/10.1016/j.semcdb.2010.09.014

Liu, C., Tan, F.C.K., Xiao, Z.-C., Dawe, G.S., 2015. Amyloid precursor protein enhances Nav1.6 sodium channel cell surface expression. J. Biol. Chem. 290, 12048–12057. https://doi.org/10.1074/jbc.M114.617092

Liu, J.-J., 2017. Regulation of dynein-dynactin-driven vesicular transport. Traffic Cph. Den. 18, 336–347. https://doi.org/10.1111/tra.12475

Maday, S., Twelvetrees, A.E., Moughamian, A.J., Holzbaur, E.L.F., 2014. Axonal transport: cargo-specific mechanisms of motility and regulation. Neuron 84, 292– 309. https://doi.org/10.1016/j.neuron.2014.10.019

Malhotra, J.D., Kazen-Gillespie, K., Hortsch, M., Isom, L.L., 2000. Sodium channel beta subunits mediate homophilic cell adhesion and recruit ankyrin to points of cell-cell contact. J. Biol. Chem. 275, 11383–11388.

Malhotra, J.D., Koopmann, M.C., Kazen-Gillespie, K.A., Fettman, N., Hortsch, M., Isom, L.L., 2002. Structural requirements for interaction of sodium channel beta 1 subunits with ankyrin. J. Biol. Chem. 277, 26681–26688. https://doi.org/10.1074/jbc.M202354200

Moughamian, A.J., Holzbaur, E.L.F., 2018. 13 - Cytoplasmic dynein dysfunction and neurodegenerative disease, in: King, S.M. (Ed.), Dyneins (Second Edition). Academic Press, pp. 286–315. https://doi.org/10.1016/B978-0-12-809470-9.00013-8

Moughamian, A.J., Holzbaur, E.L.F., 2012. Dynactin is required for transport initiation from the distal axon. Neuron 74, 331–343. https://doi.org/10.1016/j.neuron.2012.02.025

Moyon, S., Dubessy, A.L., Aigrot, M.S., Trotter, M., Huang, J.K., Dauphinot, L., Potier, M.C., Kerninon, C., Melik Parsadaniantz, S., Franklin, R.J.M., Lubetzki, C., 2015. Demyelination causes adult CNS progenitors to revert to an immature state and express immune cues that support their migration. J. Neurosci. Off. J. Soc. Neurosci. 35, 4–20. https://doi.org/10.1523/JNEUROSCI.0849-14.2015

Müller, M.J.I., Klumpp, S., Lipowsky, R., 2008. Tug-of-war as a cooperative mechanism for bidirectional cargo transport by molecular motors. Proc. Natl. Acad. Sci. U. S. A. 105, 4609–4614. https://doi.org/10.1073/pnas.0706825105

Namadurai, S., Yereddi, N.R., Cusdin, F.S., Huang, C.L.H., Chirgadze, D.Y., Jackson, A.P., 2015. A new look at sodium channel β subunits. Open Biol. 5, 140192. https://doi.org/10.1098/rsob.140192

O’Malley, H.A., Isom, L.L., 2015. Sodium channel β subunits: emerging targets in channelopathies. Annu. Rev. Physiol. 77, 481–504. https://doi.org/10.1146/annurev-physiol-021014-071846

Patton, D.E., Isom, L.L., Catterall, W.A., Goldin, A.L., 1994. The adult rat brain beta 1 subunit modifies activation and inactivation gating of multiple sodium channel alpha subunits. J. Biol. Chem. 269, 17649–17655.

Piaton, G., Aigrot, M.-S., Williams, A., Moyon, S., Tepavcevic, V., Moutkine, I., Gras, J., Matho, K.S., Schmitt, A., Soellner, H., Huber, A.B., Ravassard, P., Lubetzki, C., 2011. Class 3 semaphorins influence oligodendrocyte precursor recruitment and remyelination in adult central nervous system. Brain J. Neurol. 134, 1156–1167. https://doi.org/10.1093/brain/awr022

Rasband, M.N., 2008. Na+ channels get anchored…with a little help. J. Cell Biol. 183, 975–977. https://doi.org/10.1083/jcb.200811086

Roberts, A.J., Goodman, B.S., Reck-Peterson, S.L., 2014. Reconstitution of dynein transport to the microtubule plus end by kinesin. eLife 3, e02641.

Schroer, T.A., 2004. Dynactin. Annu. Rev. Cell Dev. Biol. 20, 759–779. https://doi.org/10.1146/annurev.cellbio.20.012103.094623

Sherman, D.L., Tait, S., Melrose, S., Johnson, R., Zonta, B., Court, F.A., Macklin, W.B., Meek, S., Smith, A.J.H., Cottrell, D.F., Brophy, P.J., 2005. Neurofascins are required to establish axonal domains for saltatory conduction. Neuron 48, 737–742. https://doi.org/10.1016/j.neuron.2005.10.019

Spassky, N., Olivier, C., Cobos, I., LeBras, B., Goujet-Zalc, C., Martínez, S., Zalc, B., Thomas, J.L., 2001. The early steps of oligodendrogenesis: insights from the study of the plp lineage in the brain of chicks and rodents. Dev. Neurosci. 23, 318–326. https://doi.org/10.1159/000048715

Stedehouder, J., Couey, J.J., Brizee, D., Hosseini, B., Slotman, J.A., Dirven, C.M.F., Shpak, G., Houtsmuller, A.B., Kushner, S.A., 2017. Fast-spiking Parvalbumin Interneurons are Frequently Myelinated in the Cerebral Cortex of Mice and Humans. Cereb. Cortex N. Y. N 1991 27, 5001–5013. https://doi.org/10.1093/cercor/bhx203

Susuki, K., Chang, K.-J., Zollinger, D.R., Liu, Y., Ogawa, Y., Eshed-Eisenbach, Y., Dours-Zimmermann, M.T., Oses-Prieto, J.A., Burlingame, A.L., Seidenbecher, C.I., Zimmermann, D.R., Oohashi, T., Peles, E., Rasband, M.N., 2013. Three mechanisms assemble central nervous system nodes of Ranvier. Neuron 78, 469–482. https://doi.org/10.1016/j.neuron.2013.03.005

Tokito, M.K., Howland, D.S., Lee, V.M., Holzbaur, E.L., 1996. Functionally distinct isoforms of dynactin are expressed in human neurons. Mol. Biol. Cell 7, 1167–1180.

Urnavicius, L., Zhang, K., Diamant, A.G., Motz, C., Schlager, M.A., Yu, M., Patel, N.A., Robinson, C.V., Carter, A.P., 2015. The structure of the dynactin complex and its interaction with dynein. Science 347, 1441–1446. https://doi.org/10.1126/science.aaa4080

Waterman-Storer, C.M., Karki, S.B., Kuznetsov, S.A., Tabb, J.S., Weiss, D.G., Langford, G.M., Holzbaur, E.L., 1997. The interaction between cytoplasmic dynein and dynactin is required for fast axonal transport. Proc. Natl. Acad. Sci. U. S. A. 94, 12180–12185.

Winters, J.J., Isom, L.L., 2016. Developmental and Regulatory Functions of Na(+) Channel Non-pore-forming β Subunits. Curr. Top. Membr. 78, 315–351. https://doi.org/10.1016/bs.ctm.2016.07.003

Wisco, D., Anderson, E.D., Chang, M.C., Norden, C., Boiko, T., Fölsch, H., Winckler, B., 2003. Uncovering multiple axonal targeting pathways in hippocampal neurons. J. Cell Biol. 162, 1317–1328. https://doi.org/10.1083/jcb.200307069

Xu, D.-E., Zhang, W.-M., Yang, Z.Z., Zhu, H.-M., Yan, K., Li, S., Bagnard, D., Dawe, G.S., Ma, Q.-H., Xiao, Z.-C., 2014. Amyloid precursor protein at node of Ranvier modulates nodal formation. Cell Adhes. Migr. 8, 396–403. https://doi.org/10.4161/cam.28802

Xue, X., Jaulin, F., Espenel, C., Kreitzer, G., 2010. PH-domain-dependent selective transport of p75 by kinesin-3 family motors in non-polarized MDCK cells. J. Cell Sci. 123, 1732–1741. https://doi.org/10.1242/jcs.056366

Yang, Y., Lacas-Gervais, S., Morest, D.K., Solimena, M., Rasband, M.N., 2004. BetaIV spectrins are essential for membrane stability and the molecular organization of nodes of Ranvier. J. Neurosci. Off. J. Soc. Neurosci. 24, 7230–7240. https://doi.org/10.1523/JNEUROSCI.2125-04.2004

Zhang, Y., Bekku, Y., Dzhashiashvili, Y., Armenti, S., Meng, X., Sasaki, Y., Milbrandt, J., Salzer, J.L., 2012. Assembly and maintenance of nodes of ranvier rely on distinct sources of proteins and targeting mechanisms. Neuron 73, 92–107. https://doi.org/10.1016/j.neuron.2011.10.016

Zonta, B., Tait, S., Melrose, S., Anderson, H., Harroch, S., Higginson, J., Sherman, D.L., Brophy, P.J., 2008. Glial and neuronal isoforms of Neurofascin have distinct roles in the assembly of nodes of Ranvier in the central nervous system. J. Cell Biol. 181, 1169–1177. https://doi.org/10.1083/jcb.200712154

