## Supplemental Figures for "An alternative mechanism of early nodal clustering and myelination onset in GABAergic neurons of the central nervous system"

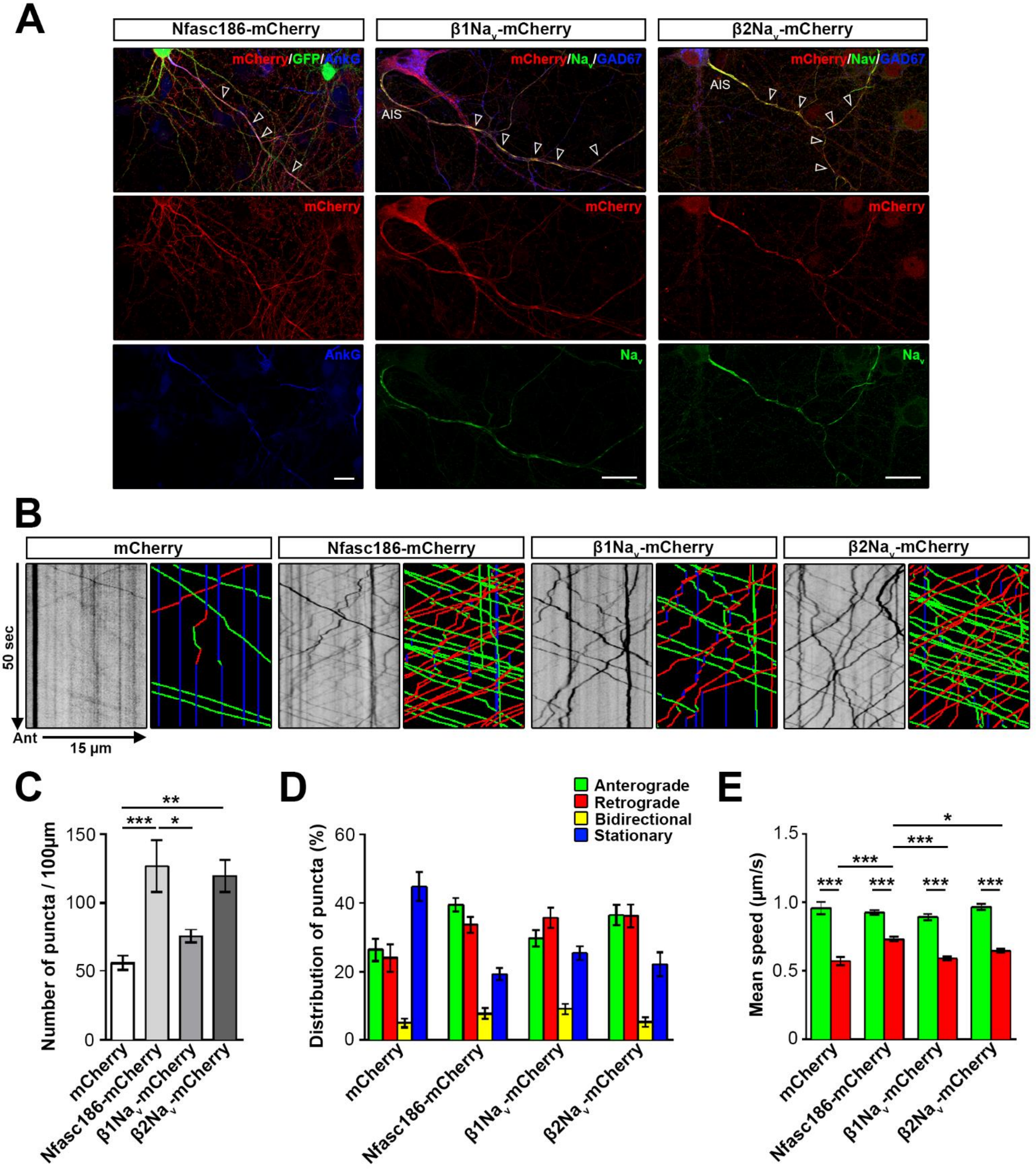


**Figure S1. Characterization of axonal trafficking of early cluster CAM markers.**
**(A)** CAM proteins tagged with fluorescent markers localize as the endogenous proteins at the axon initial segment (AIS) and early clusters in GABAergic neurons at 17 DIV. Nfasc186-mCherry (red) colocalizes with ankyrinG (AnkG, blue) at early clusters (arrowheads) in transduced VGAT-venus+ neurons (GFP, green). β1$Na_v$-mCherry and β2$Na_v$-mCherry are similarly located at both AIS and early clusters (colocalization with α$Na_v$, green, arrowheads) in GABAergic neurons (GAD67, blue). **(B)** Trafficking of early cluster CAM markers tagged with fluorescent proteins at 17 DIV. Kymographs illustrating vesicles trajectories were generated from the corresponding movies. Anterograde (Ant), retrograde, and stationary trajectories are indicated in green, red and blue respectively. Mean number of puncta **(C)**, distribution of puncta across all types of movements **(D)** and mean speed **(E)** analyzed in each condition. Typical transport includes alternating moving and pausing phases with mean velocities approaching 0.9 μm/s anterogradely and 0.6 μm/s retrogradely. Anterograde, retrograde, bidirectional movements and stationary structures are indicated in green, red, yellow and blue respectively. Data are mean ± SEM of n=17 to 21 neurons per condition from n=3 experiments. Significance was assessed by one-way analysis of variance with Tukey's post-hoc test in case of mean number of puncta and mean speed (*, $p< 0.05$; **, $p < 0.01$; and ***, $p < 0.001$) and by Chi-squared test for distribution of puncta (see Table S2 for exact *p* -values).

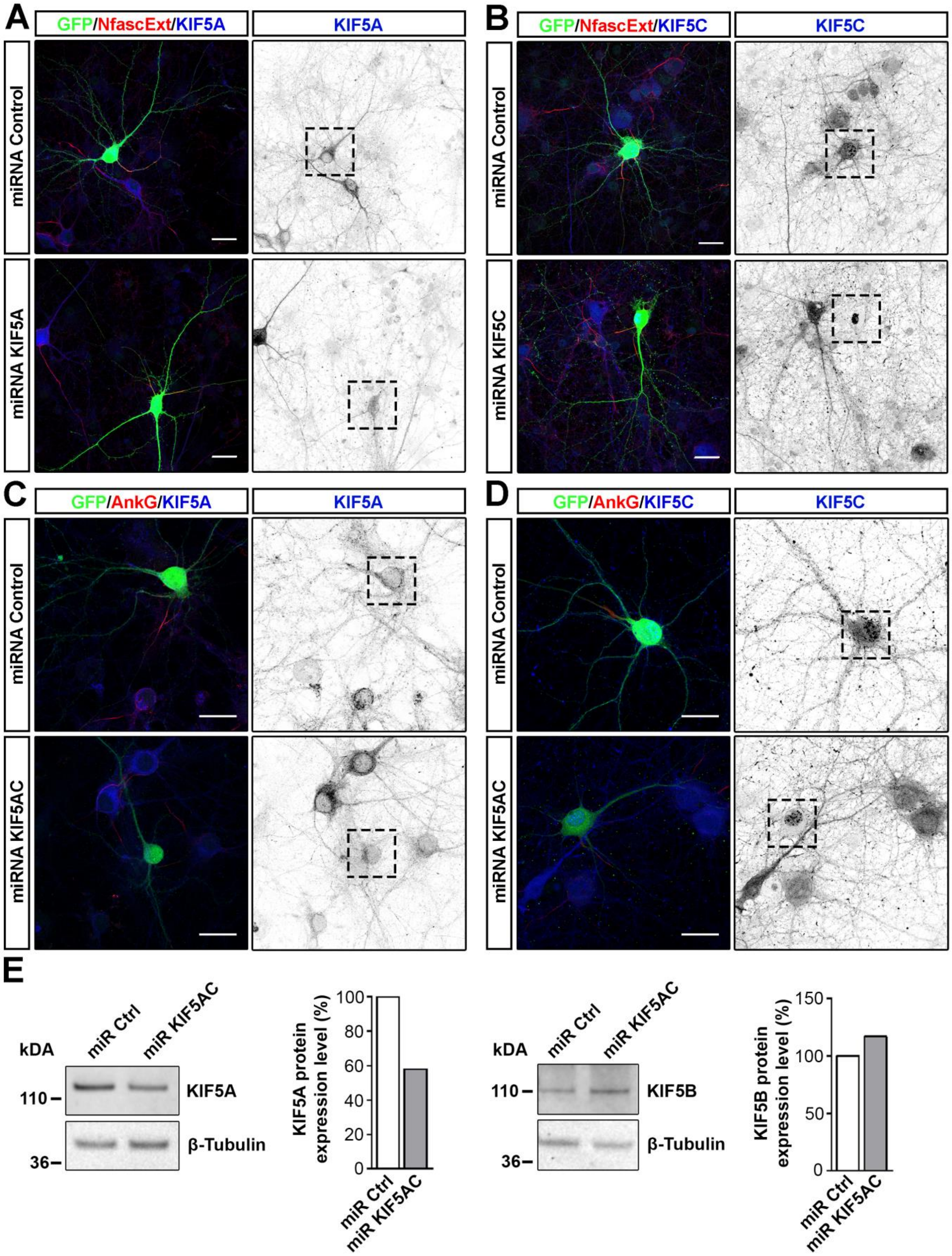


**Figure S2. Knockdown of KIF5A, KIF5C and KIF5AC in mixed hippocampal cultures.**

KIF5A (blue) expression is strongly reduced in KIF5A miRNA **(A)** and KIF5AC double-knockdown miRNA expressing neurons **(C)** compared to non-transfected and control miRNA expressing neurons. KIF5C immunolabeling (blue) is abolished in KIF5C miRNA **(B)** and KIF5AC miRNA **(D)** expressing neurons. **(E)** Protein expression levels of KIF5A and KIF5B were examined by western blotting and quantified in PC12 cells transfected with miRNA Control or miRNA KIF5AC constructs. KIF5A and KIF5B protein expression levels were normalized by β-Tubulin protein expression level. miRNA Control was considered as the reference (100%). The results were duplicated.

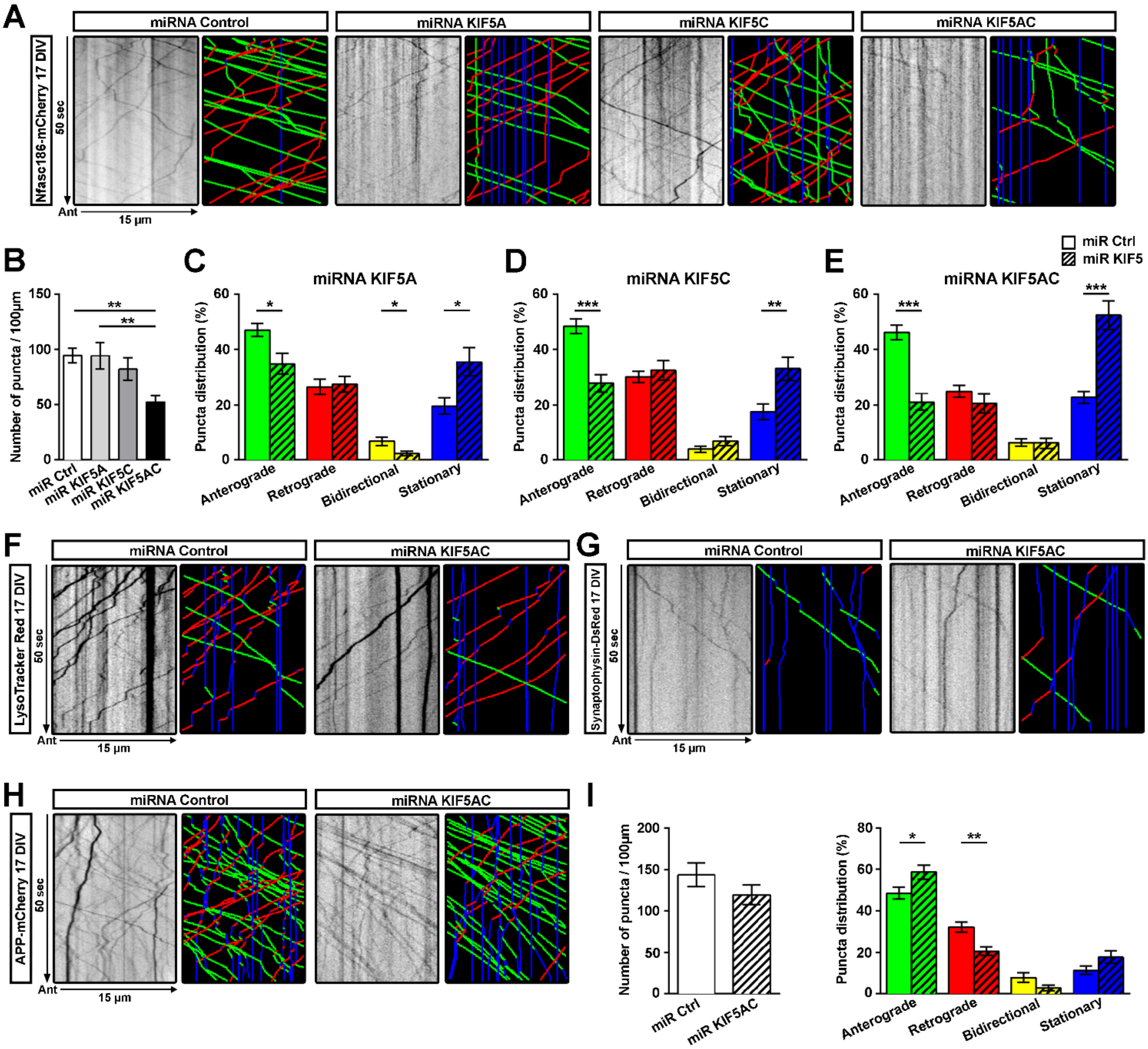


**Figure S3. Synergistic role of KIF5A and KIF5C in Nfasc186-mCherry axonal transport.**

**(A)** Kymographs and corresponding tracks of axonal Nfasc186-mCherry vesicles derived from neurons expressing miRNA Control or miRNA KIF5A, C or KIF5AC at 17 DIV. Mean number **(B)** and distribution of Nfasc186-mCherry puncta in KIF5A **(C)**, KIF5C **(D)** and KIF5AC **(E)** knock-down or in control condition. Kymographs and corresponding tracks of lysosomes **(F)**, Synaptophysin-DsRed **(G)** and APP-mCherry **(H)** transport at 17 DIV in neurons transfected with miRNA Control or miRNA KIF5AC constructs. **(I)** Mean number and distribution of APP-mCherry puncta in miRNA Control or miRNA KIF5AC conditions. Data are mean ± SEM of n=17 to 24 neurons per condition from n=3 independent experiments. One-way ANOVA with Tukey's post-hoc **(B)** and student's two-tailed unpaired t-test **(C, D, E, I)** ; *, $p < 0.05$; **, $p < 0.01$; and ***, $p < 0.001$. The difference of puncta distribution between knock-down conditions (KIF5A, C and AC) was assessed using Chi-squared test (see Table S2 for exact *p*-values).

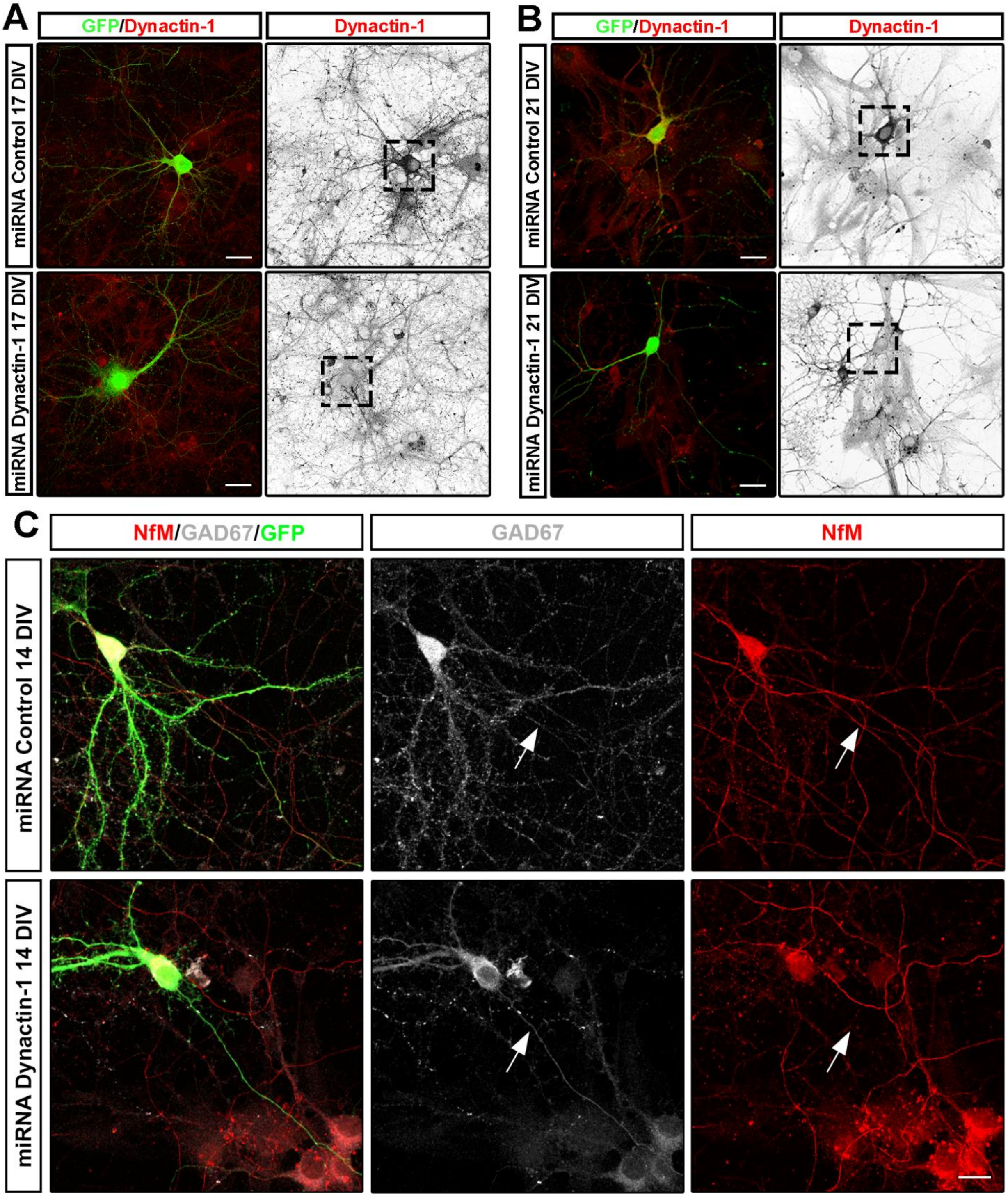


**Figure S4. Validation of loss of Dynactin-1 expression and resulting delay in maturation at 14 DIV.**
Dynactin-1 (red) expression is abolished in Dynactin-1 miRNA positive neurons at 17 DIV **(A)** and 21 DIV **(B)** compared to control conditions. miRNA positive cells are detected by GFP expression, in green, and their soma is indicated by a black box. Scale bars, 25 µm. **(C)** Neurofilament M (NfM, red, white arrow) expression is reduced in GABAergic neurons (GAD67, white) with Dynactin-1 miRNA expression (green) compared to control miRNA expressing neurons (green). Scale bar, 25 µm.

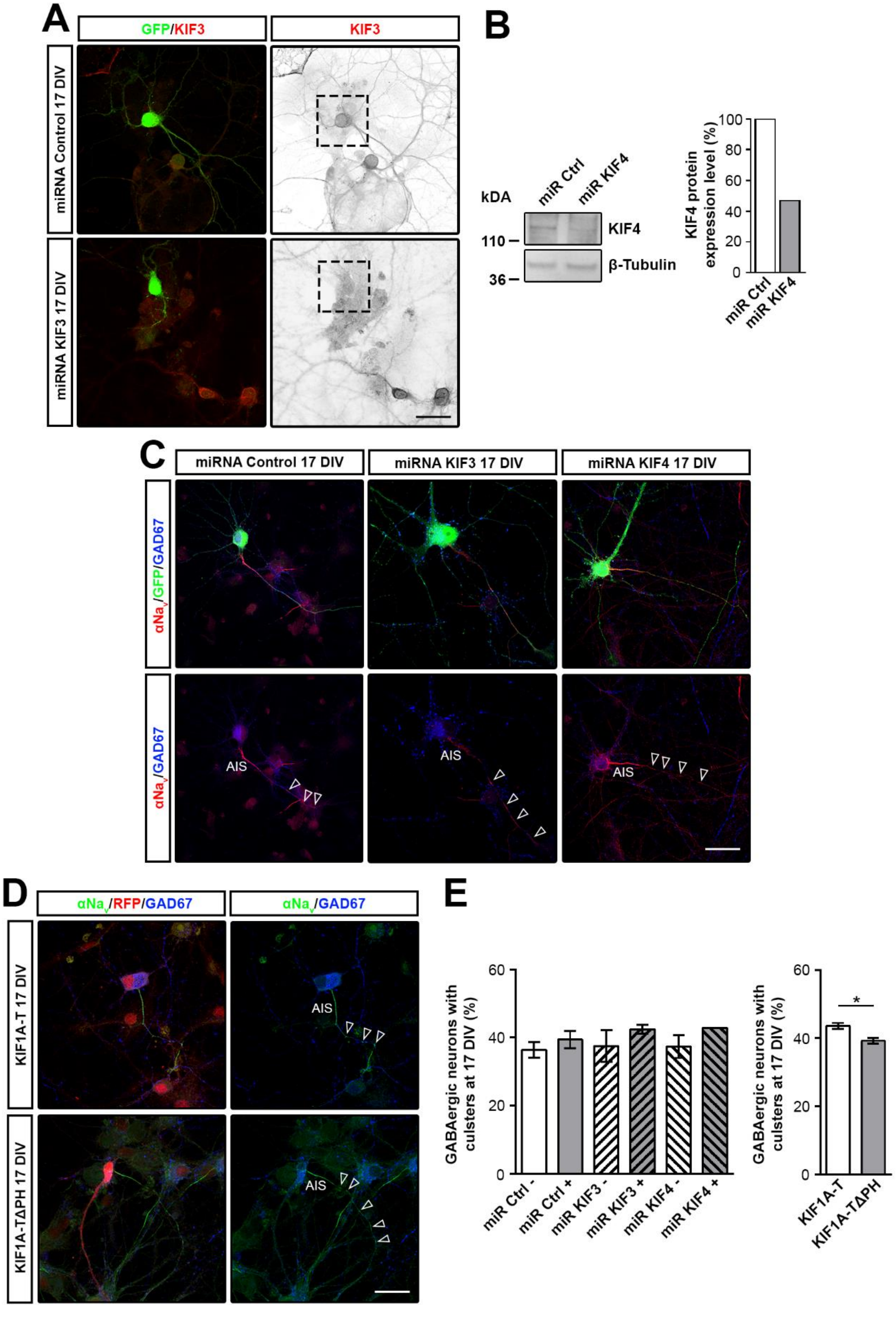


**Figure S5. KIF3, KIF4 and KIF1A loss-of-function do not impact early cluster assembly.**
**(A)** KIF3 (red) expression is strongly abolished in KIF3 miRNA expressing neurons (green) compared to non-transfected and control miRNA expressing neurons (green). **(B)** Expression level of KIF4 was determined by western blotting and quantified in N2a cells transfected with miRNA Control or miRNA KIF4 constructs. KIF4 protein expression level was normalized by β-Tubulin protein expression. miRNA Control was considered as the reference (100%). The results were duplicated. **(C)** The presence of early structures ($Na_v$, red, arrowhead) is preserved in GABAergic neurons (GAD67, blue) expressing KIF3 miRNA or KIF4 miRNA (GFP, green) compared to control condition (GFP, green) at 17 DIV. **(D)** GABAergic neurons (GAD67, blue) expressing KIF1A dominant negative (KIF1A-T, RFP, red) exhibit early clusters ($Na_v$, green, arrowheads) similarly to control condition (KIF1A-TΔPH, RFP, red) at 17 DIV. **(E)** Quantification of GABAergic neurons with clusters at 17 DIV in non-transfected neurons (miR Ctrl -/miR KIF3 -/miR KIF4-) compared to transfected neurons with a miRNA Control (miR Ctrl +), miRNA KIF3 (miR KIF3 +) or miRNA KIF4 (miR KIF4 +) expressing constructs as well as in KIF1A-T or KIF1A-TΔPH expressing neurons. Scale bars, 25 µm. AIS : axon initial segment. Mean ± SEM of 3 independent experiments. Student's two-tailed unpaired t-test; *, $p < 0.05$; **, $p < 0.01$; and ***, $p < 0.001$ (see Table S2 for exact *p*-values).

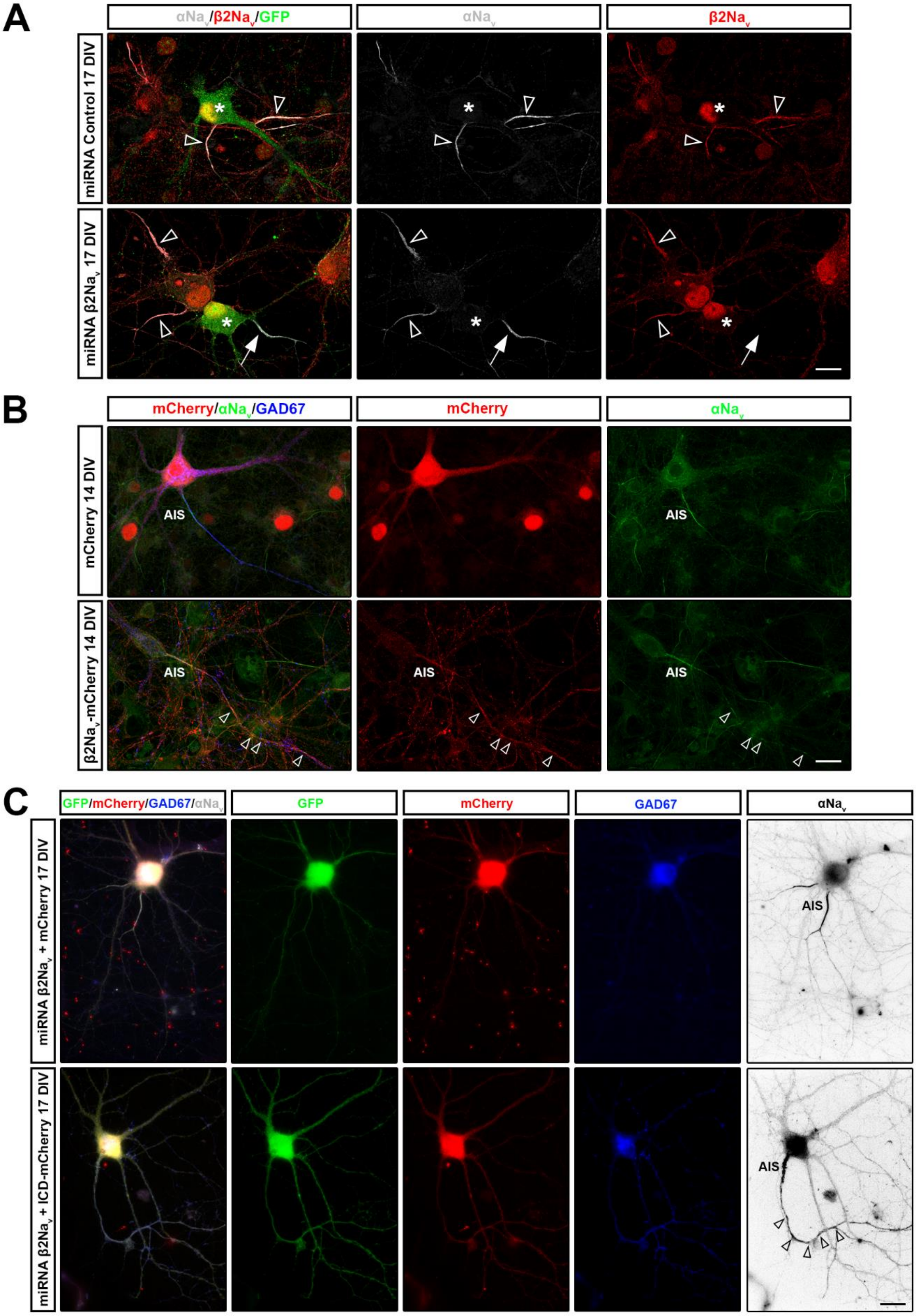


**Figure S6. β2Na$_v$ knockdown and overexpression in mixed hippocampal cultures.**

**(A)** β2Na$_v$ (red) expression is absent in axonal initial segment (AIS; white arrow) in β2Na$_v$ miRNA expressing neurons compared to non-transfected neurons and control miRNA neurons (AIS; arrowheads) while αNa$_v$ (white) expression is preserved at 17 DIV. Cell bodies of transfected neurons are indicated by an asterisk. Scale bar, 25 µm. **(B)** At 14 DIV, GABAergic neurons (GAD67, blue) overexpressing β2Na$_v$-mCherry (red) show an increased clustering (αNa$_v$, green, arrowheads) compare to neurons overexpressing mCherry (red). AIS: axon initial segment. Scale bar, 25 µm. **(C)** At 17 DIV, GABAergic neurons (GAD67, blue) transfected with the β2Na$_v$ miRNA (GFP, green) co-expressing mCherry or ICD-mCherry (red) show a partial rescue of early clustering (αNa$_v$, white) by ICD in absence of β2Na$_v$. Scale bar, 20 µm.
