## Supplemental Table 1 for "An alternative mechanism of early nodal clustering and myelination onset in GABAergic neurons of the central nervous system"

**Table S1. Oligonucleotides used for constructs generation**

| | |
|---|---|
| HindIII GFP-FW | GCAAGCTTATGGTGAGCAAGGGCGAG |
| KpnI GFP-RV | ATGGTACCTTACTTGTACAGCTCGTCCATGC |
| EcoRI b1N-FW | GTGAATTCATGGGGACGCT GGTGAATTCATGGGGACGCTGCTGGCTCT |
| HIII b1N-RV | GCAAGCTTTTCAGCCACCTGGACGCC |
| EcoRI b2N-FW | GAGGAATTCATGCACAGAGATGCCTGGC |
| HIII b2N-RV | CACAAGCTTCTTGGCGCCATCATCCGGT |
| miRNADyn-T | TGCTGTGAACTTGATGTGGTCAGCCAGTTTTGGCCACTGACTGACTGGCTGACCATCAAGTTCA |
| miRNADyn-B | CCTGTGAACTTGATGGTCAGCCAGTCAGTCAGTGGCCAAAACTGGCTGACCACATCAAGTTCAC |
| miRNAKIF5A-T | TGCTGTTAGGAAGCTCACAACGCAGAGTTTTGGCCACTGACTGACTCTGCGTTGAGCTTCCTAA |
| miRNAKIF5A-B | CCTGTTAGGAAGCTCAACGCAGAGTCAGTCAGTGGCCAAAACTCTGCGTTGTGAGCTTCCTAAC |
| miRNAKIF5C-T | TGCTGTTGAGAAGCAGGTTCAGGATCGTTTTGGCCACTGACTGACGATCCTGACTGCTTCTCAA |
| miRNAKIF5C-B | CCTGTTGAGAAGCAGTCAGGATCGTCAGTCAGTGGCCAAAACGATCCTGAACCTGCTTCTCAAC |
| miRNAScn2b-T | TGCTGTATAGCAGGAGTTGAAGGTACGTTTTGGCCACTGACTGACGTACCTTCCTCCTGCTATA |
| miRNAScn2b-B | CCTGTATAGCAGGAGGAAGGTACGTCAGTCAGTGGCCAAAACGTACCTTCAACTCCTGCTATAC |
