## Supplemental Table 2 for "An alternative mechanism of early nodal clustering and myelination onset in GABAergic neurons of the central nervous system"

**Table S2. Biostatistical analyses**

| | Test used | | n | | | Descriptive stats (average, variance) | | P value | |
|---|---|---|---|---|---|---|---|---|---|
| Figure number | Which test ? | Section & paragraph | Exact value | Defined ? | Section & paragraph | Reported ? | Section & Paragraph | Exact value | Section & Paragraph |
| Fig. S1 C | One- way ANOVA | Fig. legend & Methods | **17** (mCh), **21** (Nfasc186-mCh), **18** (β1Nav-mCh), **18** (β2Nav-mCh) neurons | 3 independent embryonic cultures | Fig. legend | Mean ± SEM | Fig. legend | Number of puncta **: Grouped analysis** : p = 0.0002 ; **Multiple comparisons** : p = 0.0008 (mCh vs. Nfasc186-mCh) ; p = 0.0047 (mCh vs. β2Nav-mCh) ; p = 0.0220 (Nfasc186-mCh vs. β1Nav-mCh) | Results; Paragraph 1 |
| Fig. S1 D | Pearson's Chi-squared test | Fig. legend & Methods | **17** (mCh), **21** (Nfasc186-mCh), **18** (β1Nav-mCh), **18** (β2Nav-mCh) neurons | 3 independent embryonic cultures | Fig. legend | Mean ± SEM | Fig. legend | Puncta distribution : p = 0.0497 | Results; Paragraph 1 |
| Fig. S1 E | One- way ANOVA | Fig. legend & Methods | **17** (mCh), **21** (Nfasc186-mCh), **18** (β1Nav-mCh), **18** (β2Nav-mCh) neurons | 3 independent embryonic cultures | Fig. legend | Mean ± SEM | Fig. legend | Mean Speed **: Grouped analysis** : p < 0.0001; **Multiple comparisons** : p < 0.0001 (mCh Ant vs. Ret; Nfasc186-mCh Ant vs. Ret; β1Nav-mCh Ant vs. Ret; β2Nav-mCh Ant vs. Ret ; mCh Ret vs. Nfasc186-mCh Ret; Nfasc186-mCh Ret vs β1Nav-mCh Ret) ; p = 0.0152 (Nfasc186-mCh Ret vs β2Nav-mCh Ret) | Results; Paragraph 1 |
| Fig. 1 B | One- way ANOVA | Fig. legend & Methods | **18** (β2Nav-GFP+Nfasc186-mCh), **22** (β2Nav-GFP+β1Nav-mCh), **17** (β2Nav-GFP+APP-mCh) neurons | 3 independent embryonic cultures | Fig. legend | Mean ± SEM | Fig. legend | Number of puncta **: Grouped analysis** : p = 0.0465 ; **Multiple comparisons** : p = 0.0359 (β2Nav-GFP+β1Nav-mCh vs β2Nav-GFP+APP-mCh) | Results; Paragraph 2 |
| Fig. 1 C | One- way ANOVA | Fig. legend & Methods | **18** (β2Nav-GFP+Nfasc186-mCh), **22** (β2Nav-GFP+β1Nav-mCh), **17** (β2Nav-GFP+APP-mCh) neurons | 3 independent embryonic cultures | Fig. legend | Mean ± SEM | Fig. legend | **Distribution of co-labeled puncta : Grouped analysis** : p = 0.6365 | Results; Paragraph 2 |
| Fig. 1 D | One- way ANOVA | Fig. legend & Methods | **18** (β2Nav-GFP+Nfasc186-mCh), **22** (β2Nav-GFP+β1Nav-mCh), **17** (β2Nav-GFP+APP-mCh) neurons | 3 independent embryonic cultures | Fig. legend | Mean ± SEM | Fig. legend | **Percentage of co-labeled puncta : Grouped analysis** : p < 0.0001; **Multiple comparisons** : **Ant puncta** : p < 0.0001 (β2Nav-GFP+β1Nav-mCh vs. β2Nav-GFP+APP-mCh ; β2Nav-GFP+Nfasc186-mCh vs. β2Nav-GFP+APP-mCh); **Ret puncta** : p < 0.0001 (β2Nav-GFP+Nfasc186-mCh vs. β2Nav-GFP+APP-mCh) | Results; Paragraph 2 |
| Fig. 1 D | Pearson's Chi-squared test | Fig. legend & Methods | **18** (β2Nav-GFP+Nfasc186-mCh), **22** (β2Nav-GFP+β1Nav-mCh), **17** (β2Nav-GFP+APP-mCh) neurons | 3 independent embryonic cultures | Fig. legend | Mean ± SEM | Fig. legend | **Puncta distribution :** p = 0.0035 | Results; Paragraph 2 |
| Fig. S3 B | One- way ANOVA | Fig. legend & Methods | **20** (miR Ctrl), **22** (miR KIF5A), **22** (miR KIF5C), **24** (miR KIF5AC) neurons | 3 independent embryonic cultures | Fig. legend | Mean ± SEM | Fig. legend | Number of puncta **: Grouped analysis** : p = 0.0027 ; **Multiple comparisons** : p = 0.0150 (miR Ctrl vs miR KIF5AC) ; p = 0.0073 (miR KIF5A vs miR KIF5AC) | Results; Paragraph 3 |
| Fig. S3 C | Unpaired *t* -test | Fig. legend & Methods | **17** (miR Ctrl), **22** (miR KIF5A) neurons | 3 independent embryonic cultures | Fig. legend | Mean ± SEM | Fig. legend | **Ant puncta** : p = 0.0141 (miR Ctrl vs miR KIF5A) ; **Bi puncta** : p = 0.0126 (miR Ctrl vs miR KIF5A) ; **St puncta** : p = 0.0160 (miR Ctrl vs miR KIF5A) | Results; Paragraph 3 |
| Fig. S3 D | Unpaired *t* -test | Fig. legend & Methods | **18** (miR Ctrl), **22** (miR KIF5C) neurons | 3 independent embryonic cultures | Fig. legend | Mean ± SEM | Fig. legend | **Ant puncta** : p < 0.0001 (miR Ctrl vs miR KIF5C) ; **St puncta** : p = 0.0052 (miR Ctrl vs miR KIF5C) | Results; Paragraph 3 |
| Fig. S3 E | Unpaired *t* -test | Fig. legend & Methods | **20** (miR Ctrl), **24** (miR KIF5AC) neurons | 3 independent embryonic cultures | Fig. legend | Mean ± SEM | Fig. legend | **Ant puncta** : p < 0.0001 (miR Ctrl vs miR KIF5AC) ; **St puncta** : p < 0.0001 (miR Ctrl vs miR KIF5AC) | Results; Paragraph 3 |
| Fig. S3 I | Unpaired t-test | Fig. legend & Methods | **22** (miR Ctrl), **18** (miR KIF5AC) neurons | 4 independent embryonic cultures | Fig. legend | Mean ± SEM | Fig. legend | **Number of puncta** : p = 0.2077 (miR Ctrl vs miR KIF5AC) ; **Ant puncta** : p = 0.0252 (miR Ctrl vs miR KIF5AC) ; **Ret puncta** : p = 0.0012 (miR Ctrl vs miR KIF5AC) | Results; Paragraph 3 |
| Fig. S3 C, D, I | Pearson's Chi-squared test | Fig. legend & Methods | **17 to 22** (miR Ctrl), **22** (miR KIF5A), **22** (miR KIF5C) and **18** (miR KIF5AC) neurons | 3 independent embryonic cultures | Fig. legend | Mean ± SEM | Fig. legend | **Puncta distribution** : p = 0.0401 (miR Ctrl vs miR KIF5A of Nfasc186-mCh) ; p = 0.0124 (miR Ctrl vs miR KIF5C of Nfasc186-mCh) ; p = 0.1302 (miR Ctrl vs miR KIF5AC of APP-mCh DIV17) | Results; Paragraph 3 |
| Fig. 2 B | Unpaired *t* -test | Fig. legend & Methods | **20** (miR Ctrl), **24** (miR KIF5AC) neurons | 3 independent embryonic cultures | Fig. legend | Mean ± SEM | Fig. legend | **Number of puncta :** p < 0.0001 (miR Ctrl vs miR KIF5AC) ; **Ant puncta** : p < 0.0001 (miR Ctrl vs miR KIF5AC) ; **St puncta** : p < 0.0001 (miR Ctrl vs miR KIF5AC) | Results; Paragraph 3 |
| Fig. 2 D | Unpaired *t* -test | Fig. legend & Methods | **20** (miR Ctrl), **21** (miR KIF5AC) neurons | 3 independent embryonic cultures | Fig. legend | Mean ± SEM | Fig. legend | **Number of puncta :** p < 0.0001 (miR Ctrl vs miR KIF5AC) ; **Ant puncta** : p = 0.0026 (miR Ctrl vs miR KIF5AC) ; **Ret puncta** : p = 0.0202 (miR Ctrl vs miR KIF5AC) ; **Bi puncta** : p = 0.0440 (miR Ctrl vs miR KIF5AC) ; **St puncta** : p = 0.0002 (miR Ctrl vs miR | Results; Paragraph 3 |
| Fig. 2 F | Unpaired *t* -test | Fig. legend & Methods | **21** (miR Ctrl), **21** (miR KIF5AC) neurons | 3 independent embryonic cultures | Fig. legend | Mean ± SEM | Fig. legend | **Number of puncta :** p < 0.0001 (miR Ctrl vs miR KIF5AC) ; **Ret puncta** : p = 0.0005 (miR Ctrl vs miR KIF5AC) ; **St puncta** : p < 0.0001 (miR Ctrl vs miR KIF5AC) | Results; Paragraph 3 |
| Fig. 2 H | Unpaired *t* -test | Fig. legend & Methods | **21** (miR Ctrl), **21** (miR KIF5AC) neurons | 3 independent embryonic cultures | Fig. legend | Mean ± SEM | Fig. legend | **Number of puncta :** p < 0.0001 (miR Ctrl vs miR KIF5AC) ; **Ant puncta** : p = 0.0004 (miR Ctrl vs miR KIF5AC) ; **Ret puncta** : p = 0.0002 (miR Ctrl vs miR KIF5AC) ; **St puncta** : p < 0.0001 (miR Ctrl vs miR KIF5AC) | Results; Paragraph 3 |
| Fig. 2 B, D, F, H | Pearson's Chi-squared test | Fig. legend & Methods | **20** (miR Ctrl) to **24** (miR KIF5AC) neurons | 3 independent embryonic cultures | Fig. legend | Mean ± SEM | Fig. legend | **Puncta distribution** : p = 0.0055 (miR Ctrl vs miR KIF5AC of β1Nav-mCh) ; p = 0.0020 (miR Ctrl vs miR KIF5AC of Nfasc186-mCh) ; p = 0.0045 (miR Ctrl vs miR KIF5AC of β2Nav-mCh DIV17) ; p = 0.0005 (miR Ctrl vs miR KIF5AC of β2Nav-mCh DIV10) | Results; Paragraph 3 |
| Fig. 3 B | Unpaired *t* -test | Fig. legend & Methods | **24** (miR Ctrl), **25** (miR Dyn-1) neurons | 3 independent embryonic cultures | Fig. legend | Mean ± SEM | Fig. legend | **Number of puncta :** p = 0.0006 (miR Ctrl vs miR Dyn-1) ; **Ret puncta** : p < 0.0021 (miR Ctrl vs miR Dyn-1) ; **St puncta** : p < 0.0001 (miR Ctrl vs miR Dyn-1) | Results; Paragraph 4 |
| Fig. 3 D | Unpaired *t* -test | Fig. legend & Methods | **18** (miR Ctrl), **20** (miR Dyn-1) neurons | 3 independent embryonic cultures | Fig. legend | Mean ± SEM | Fig. legend | **Number of puncta :** p = 0.0872 (miR Ctrl vs miR Dyn-1) ; **Ret puncta** : p = 0.0035 (miR Ctrl vs miR Dyn-1) ; **St puncta** : p = 0.0042 (miR Ctrl vs miR Dyn-1) | Results; Paragraph 4 |
| Fig. 3 F | Unpaired *t* -test | Fig. legend & Methods | **19** (miR Ctrl), **19** (miR Dyn-1) neurons | 3 independent embryonic cultures | Fig. legend | Mean ± SEM | Fig. legend | **Number of puncta :** p = 0.5630 (miR Ctrl vs miR Dyn-1) ; **St Puncta** : p = 0.0059 (miR Ctrl vs miR Dyn-1) | Results; Paragraph 4 |
| Fig. 3 H | Unpaired *t* -test | Fig. legend & Methods | **20** (miR Ctrl), **20** (miR Dyn-1) neurons | 3 independent embryonic cultures | Fig. legend | Mean ± SEM | Fig. legend | **Number of puncta :** p < 0,0001 (miR Ctrl vs miR Dyn-1) ; **Ant puncta** : p = 0,0019 (miR Ctrl vs miR Dyn-1) ; **Ret puncta** : p = 0.0176 (miR Ctrl vs miR Dyn-1) ; **St puncta** : p = 0.0012 (miR Ctrl vs miR Dyn-1) | Results; Paragraph 4 |
| Fig. 3 B, D, F, H | Pearson's Chi-squared test | Fig. legend & Methods | **24** (miR Ctrl) to **25** (miR Dyn-1) neurons | 3 independent embryonic cultures | Fig. legend | Mean ± SEM | Fig. legend | **Puncta distribution** : p = 0.0450 (miR Ctrl vs miR Dyn-1 of β1Nav-mCh) ; p = 0.0050 (miR Ctrl vs miR Dyn-1 of Nfasc186-mCh) ; p = 0.1228 (miR Ctrl vs miR Dyn-1 of β2Nav-mCh DIV17) ; p = 0.0002 (miR Ctrl vs miR Dyn-1 of β2Nav-mCh DIV10) | Results; Paragraph 4 |
| Fig. 4 B | Unpaired t-test | Fig. legend & Methods | At least **150** (miR Ctrl), **151** (miR KIF5AC) cells per condition, per experiment | 4 independent embryonic cultures | Fig. legend | Mean ± SEM | Fig. legend | **Percentage of GABA cells with node-like clusters** : p = 0.0020 (miR Ctrl- vs miR KIF5AC+) ; p = 0.0003 (miR KIF5AC- vs miR KIF5AC+) | Results; Paragraph 5 |
| Fig. 4 D | Unpaired t-test | Fig. legend & Methods | At least **165** (miR Ctrl), **137** (miR Dyn-1) cells per condition, per experiment | 3 independent embryonic cultures | Fig. legend | Mean ± SEM | Fig. legend | **Percentage of GABA cells with node-like clusters** : p = 0.0010 (miR Ctrl- vs miR Dyn+) ; p = 0.0022 (miR Ctrl+ vs miR Dyn+) ; p = 0.0010 (miR Dyn- vs miR Dyn+) | Results; Paragraph 5 |
| Fig. 4 F | Unpaired t-test | Fig. legend & Methods | At least **187** (miR Ctrl), **247** (miR Dyn-1) cells per condition, per experiment | 3 independent embryonic cultures | Fig. legend | Mean ± SEM | Fig. legend | **Percentage of GABA cells with node-like clusters** : p = 0.6820 (miR Ctrl- vs miR Ctrl+) ; p = 0.3078 (miR Ctrl- vs miR Dyn-) ; p = 0.8783 (miR Ctrl+ vs miR Dyn+) ; p = 0.9759 (miR Dyn- vs miR Dyn+) | Results; Paragraph 5 |
| Fig. S5 C | Unpaired t-test | Fig. legend & Methods | At least **159** (miR Ctrl), **124** (miR KIF3), **66** (miR KIF4), **9** (KIF1A-T), **8** (KIF1A-TΔPH) cells per condition, per experiment | 3 independent embryonic cultures | Fig. legend | Mean ± SEM | Fig. legend | **Percentage of GABA cells with node-like clusters** : p = 0.4199 (miR Ctrl- vs miR Ctrl+) ; p = 0.8354 (miR Ctrl- vs miR KIF3-) ; p = 0.3508 (miR Ctrl+ vs miR KIF3+) ; p = 0.3681 (miR KIF3- vs miR KIF3+) ; p = 0.5812 (miR Ctrl- vs miR KIF4-) ; p = 0.2470 (miR Ctrl+ vs miR KIF4+) ; p = 0.0937 (miR KIF4- vs miR KIF4+) ; p = 0.0250 (KIF1A-T vs KIF1A-TΔPH) | Results; Paragraph 5 |
| Fig. S5 E | Unpaired t-test | Fig. legend & Methods | **30** (KIF1A-T) and **33** (KIF1A-TΔPH) neurons | 3 independent embryonic cultures | Fig. legend | Mean ± SEM | Fig. legend | **Percentage of GABA cells with node-like clusters** : p = 0.0250 (KIF1A-T vs KIF1A-TΔPH) | Results; Paragraph 5 |
| Fig. 5 H | Unpaired t-test | Fig. legend & Methods | At least **148** (miR Ctrl), **149** (miR β2Nav), **11** (miR β2Nav + mCh), **16** (miR β2Nav + ICD-mCh), **20** (mCh), **47** (β2Nav-mCh) cells per condition, per experiment | 3 independent embryonic cultures | Fig. legend | Mean ± SEM | Fig. legend | **Percentage of GABA cells with node-like clusters** : p = 0.0175 (miR Ctrl+ vs miR β2Nav+) ; p = 0.0332 (miR β2Nav- vs miR β2Nav+) ; p = 0.0370 (miR Ctrl- vs miR β2Nav+); p = 0.0213 (miR β2Nav + mCh vs miR β2Nav + ICD-mCh); p = 0.0017 (mCh vs β2Nav) | Results; Paragraph 6 |
| Fig. 6 D | Mann-Withney | Fig. legend & Methods | **9** myelin sheaths not facing a node-like clusters and **9** myelin sheaths facing a node-like cluster | 18 independent embryonic cultures | Fig. legend | Mean ± SEM | Fig. legend | **Mean speed of myelin progression when facing or not a node-like cluster** : p = 0.0028 | Results; Paragraph 8 |
| Fig. 6 D | Mann-Withney | Fig. legend & Methods | **5** myelin sheaths facing a node-like clusters | 5 independent embryonic cultures | Fig. legend | Mean ± SEM | Fig. legend | **Mean speed of myelin progression depending on the distance from the node-like cluster (>5µm vs <5µm away from the cluster)** : p = 0.0556 | Results; Paragraph 8 |

**Legends :**

Ant = Anterograde
Ret = Retrograde
Bi = Bidirectional
St = Stationary
